# Comparative multi-OMICS single cell atlas of five COVID-19 (rAdVV and mRNA) vaccines describe unique and distinct mechanisms of action

**DOI:** 10.1101/2022.09.12.507666

**Authors:** Yogesh Singh, Antje Schulze Selting, Gisela Gabernet, Urvi Ray, Rimpi Bajaj, Mohammed Ali, Marion Loitz, Vincent Hammer, Elena Buena-Atienza, Christoph Ruschil, Jeannette Huebener-Schmid, Markus Kowarik, Madhuri S Salker, Nicolas Casadei, Sven Nahnsen, Peter Kremsner, Stephan Ossowski, Daniel M Altmann, Deutsche COVID-19 OMICS Initiative (DeCOI), Olaf Riess

## Abstract

COVID-19 vaccines based on a range of expression platforms have shown considerable protective efficacy, generating antibody and T cell immune responses. However, molecular pathways underpinning COVID-19 vaccine priming of immunity against the SARS-CoV-2 virus have not yet been explored extensively. This analysis is critical to optimization of future vaccination strategies, schedules, and combinations. Thus, we investigated a cohort of individuals pre- and post-vaccination to understand the humoral and cellular immune response against different COVID-19 vaccines, including recombinant adenoviral vector (rAdVV) and mRNA-based vaccines. Single-cell RNA sequencing allowed characterization of monocytes, T, NK and B cell activation at the transcriptomics/proteomic level, in response to different COVID-19 vaccines. Our data revealed that different COVID-19 vaccines elicit a unique and distinct mechanism of action. Specifically, we revealed that rAdVV vaccines negatively regulate CD4^+^ T cell activation, leukocytes chemotaxis, IL-18 signalling and antigen presentation by monocytes whilst mRNA vaccines positively regulate NKT cell activation, platelets activation and chemokine signalling pathways. An antigen-specific T cell response was already observed following the 1^st^ vaccine dose and was not further augmented after the subsequent 2^nd^ dose of the same vaccine and it was dependent on the type of vaccination used. Our integrated three layered-analyses highlights that COVID-19 vaccines evoke a strong but divergent immune response at the RNA, protein, and cellular levels. Our approach is able to pinpoint efficacy and mechanisms controlling immunity to vaccination and open the door for better vaccination which could induce innate and adaptive immunity equally in the long term.

**Key findings:**

1. Decrease in major three cell types classical and non-classical monocytes and NK type III cells after COVID-19 vaccination
2. Individual vaccination (AZ, JJ, MD, PB) has differential effect on various immune cell subsets and regulates unique cell populations, whilst no change was observed for CV vaccination
3. rAdVV and mRNA vaccines have different mechanism of action for activation of lymphocytes and monocytes, respectively
4. rAdVV vaccines negatively regulates CD4^+^ T cell activation, leukocytes chemotaxis, IL-18 signalling and antigen presentation whilst mRNA vaccines positively regulate NKT cell activation, platelets activation and chemokine signalling pathways.
5. An antigen-specific T cell response was prompted after the 1^st^ vaccine dose and not augmented after the subsequent 2^nd^ dose of the same vaccine.

**Graphical abstract:** 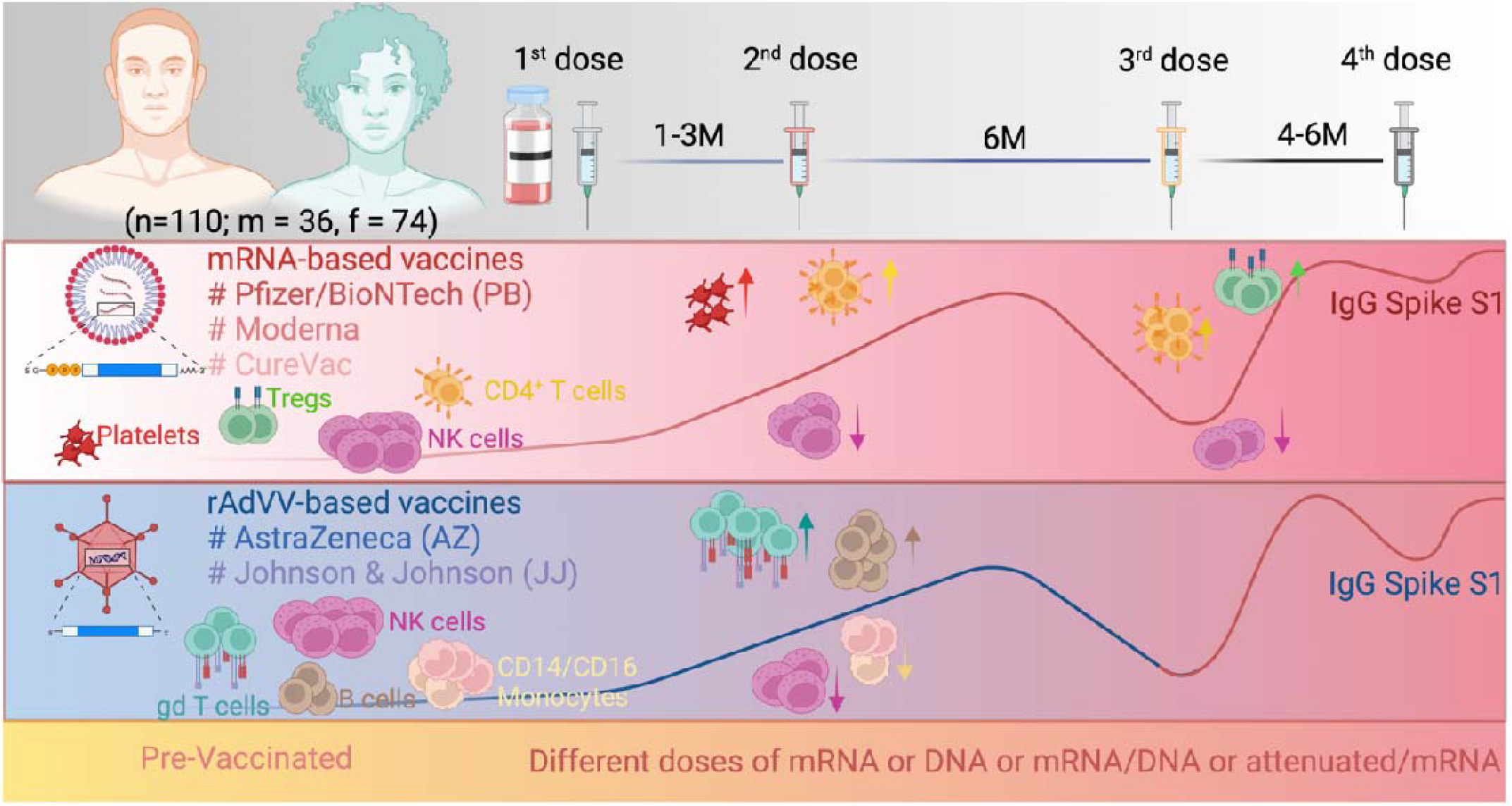

## Introduction

Vaccine development for the SARS-CoV-2 virus started shortly after the first infections were reported in October 2019. Besides conventional recombinant adenovirus vector (rAdVV)-based vaccines [ChAdOx1 nCoV-19; AstraZeneca (AZ), Ad26.CoV2.S; Johnson & Johnson or Janssen (JJ)], inactivated virus-based vaccine (BBIBP-CorV; Sinopharma (SP) COVID-19 vaccine), protein-based vaccine (NVX-CoV2373; Novavax (NV)) pharmaceutical companies for the first time employed mRNA-based vaccination technology [BNT162b1; Pfizer/BioNTech (PB), mRNA-1273; Moderna (MD), CVnCV; CureVac (CV)]^1–10^. Clinical Phase I/II trials of COVID-19 mRNA (PB) and rAdVV-based vaccines (AZ) were reported in mid-August 2020^11,12^ and between December 2020 and early 2021 several COVID-19 vaccines were approved by the regulatory agencies^13,14^. Both rAdVV and mRNA based COVID-19 vaccines against SARS-CoV-2 turned out to be crucial and effective in limiting COVID-19 severity and spread^15,16^. However, several studies showed relatively rapid waning of protective immunity over subsequent months even after booster dose^17–19^. Moreover, many individuals obtained different vaccine brand for the first and second vaccination and at the same time infection with emerging SARS-CoV-2 variants started to arise^20–23^, resulting in a complex interplay of different vaccines and SARS-CoV-2 variants.

In the past two years, several longitudinal studies have offered insights into the immune mechanisms following SARS-CoV-2 infection and/or vaccination, applying single-cell genomics, mass cytometry and flow cytometry methods^24–27^. The notion is that higher granularity analysis of molecular programs may illuminate issues such as difference in immunogenicity, efficacy, immune subsets and durability. There remains need to characterize the innate and adaptive immune activation after the second vaccine dose, and how the 3^rd^ or 4^th^ dose boosters helps to protect against new variants of concerns (VOCs). Recent studies suggest that omicron escapes the majority of existing SARS-CoV-2 neutralizing antibodies induced by vaccination or previous infections^28–30^. Thus far there has been little direct comparative analysis of how different vaccine formulations prime specific immunity considering that spike as presented in the context of either mRNA or modified adenovirus platforms is detected by the host using different dendritic cells (DCs) recognition pathways: for example, the greater importance of TLR7 in the case of mRNA vaccines encoding the SARS-CoV-2 spike (S) protein encapsulated in lipid nanoparticles, and of TLR9 in rAdVV vectors encoding the S protein^31^. Therefore, precise understanding of the immune response in vaccinated individuals is crucial to determine the clinical need for future booster vaccination programmes and to monitor immune response.

We thus performed a longitudinal study from rAdVV (AZ and JJ) and mRNA (MD, CV, PB) vaccination volunteers using multi-OMICS single cell (transcriptomics, proteogenomics) analysis and detection of T cell specific protein markers, SARS-CoV-2-specific antibodies, and cytokine levels. We monitored the evolution of the innate and adaptive immune response induced by COVID-19 vaccines from Pre-Vac (PrV) status up to the 1^st^ - 4^th^ doses. We identified that individual vaccines (rAdVV and mRNA) have a unique and distinct mechanisms of T cells activation and antigen presentation by monocytes/DCs which could alter the outcome of vaccines efficacy.

## Results

### Demographics of vaccinated individuals

We recruited in total 110 individuals (Suppl. Table 1). The overall study plan is described in Fig. 1a. We verified that none of the individuals had been previously exposed to the SARS-CoV-2 virus before by nucleocapsid (N) antibody test (discovery cohort only). Individuals positive for nucleocapsid (N) antibody test were excluded from further analyses. We split the individuals into two different cohorts. The first cohort consisted of 28 unvaccinated (naïve) and vaccinated individuals (discovery cohort; n=3-5/vaccine). PBMCs samples from this cohort were used for single cell RNA-sequencing (scRNA-seq) analysis and flow cytometry-based methods to understand the molecular, cellular, and humoral immune response to vaccination. Samples were collected before vaccination (PrV1, n=23) and after either the 1^st^ (PoV1) or the 2^nd^ dose (PoV2) of 5 different COVID-19 vaccines: AZ or mixed AZ and PB (n=4), JJ (n=3), PB (n=15), MD (n=3), CV (n=3). JJ vaccinated patients received only a single dose during this period which was considered equivalent to the two doses of other COVID-19 vaccines as recommended by regulatory authorities to use.

**Fig 1.**
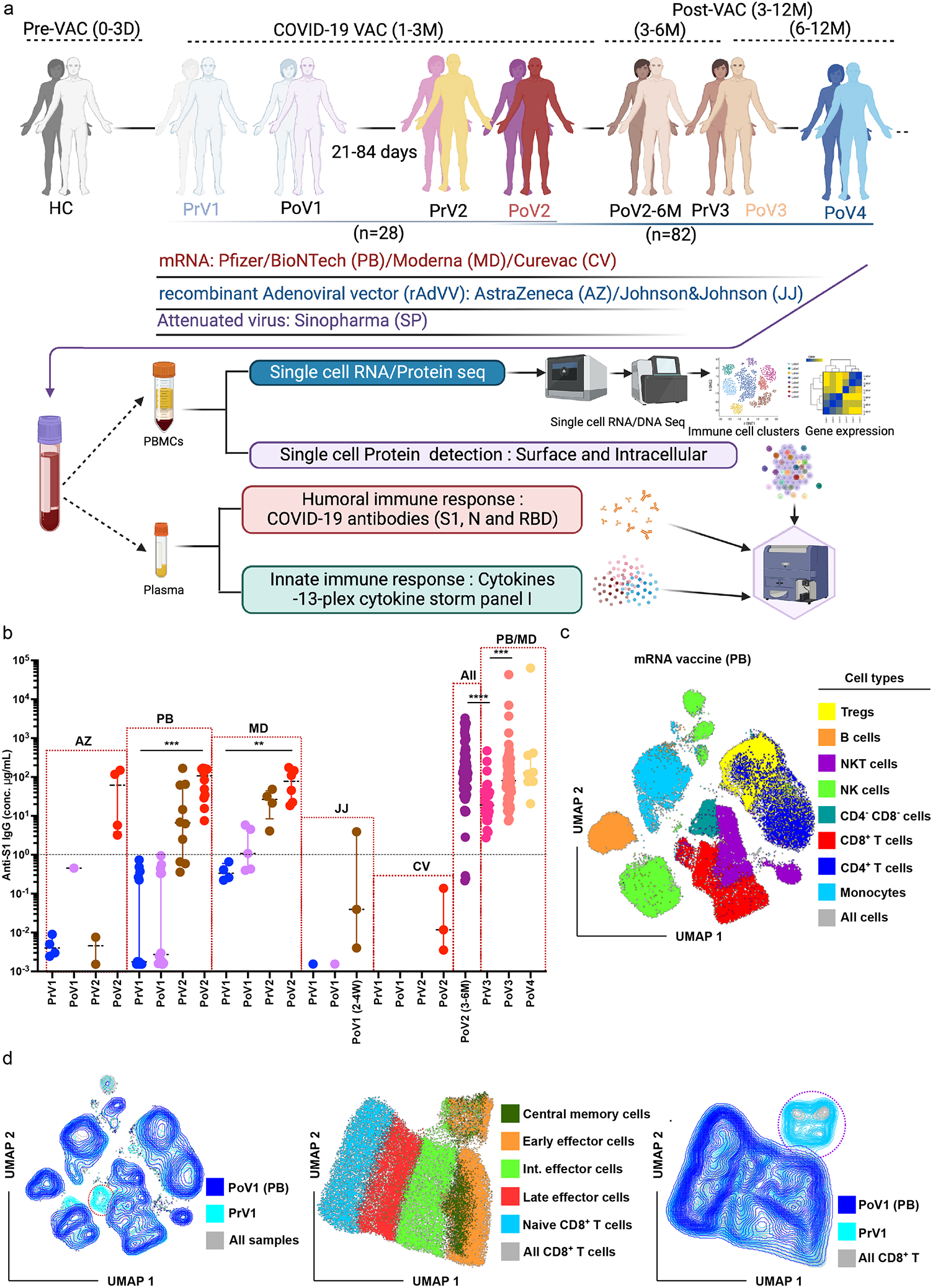
Immunological changes after SARS-CoV-2 first dose of COVID-19 vaccination. (**a**) Experimental design for the study. Blood was collected from naive individuals before vaccination (PrV1), 7-10 days after dose 1 (PoV1), and 21-84 days after the first dose and before 0-3 days for second dose (PoV2), 7-10 days after dose 2, 3-6 months (antibody and cytokine measurements), after 6 months before dose 3 (PrV3), 7-10 days after dose 3 (PoV3) and some individual 3 months post V3 and 7-10 days after dose 4 (PoV4). Immune phenotypes of PBMCs and antigen-specific T and B cell responses were measured using multicolour flow cytometry. Longitudinal serological responses to the vaccine were measured using multiplex flow cytometry-based assay for the antibodies and cytokines. PBMC samples (PrV1, PoV2, and PoV3 were profiled using multi-OMICS single cell analyses. (**b**) IgG Spike S1 antibody levels PrV1 to PoV4 vaccinated individuals. Spike S1 antibody levels were significantly upregulated after 2^nd^ dose in PB and MD COVID-19 vaccines (Kruskal Wallis test and post-hoc Dunn’s test). No statistically significant difference was observed at antibody level among 3-6 months vaccinated individuals with different vaccines, although PB had a higher amount compared with MD or mixed vaccination (AZ+PB/MD) (Kruskal Wallis test and post-hoc Dunn’s test). A group comparison among PoV2 (3-6M), PrV3, PoV3 (6-8M) and PoV4 samples, a statistical significantly (Kruskal Wallis test and post-hoc Dunn’s test) increased Spike S1 antibody levels was reached for PoV2(3-6M) vs PrV3 (p<0.0001), PrV3 vs PoV3 (p=0.0008). Error bars denote medians and interquartile ranges. \**P* < 0.05; \*\**P* < 0.01; \*\*\**P* < 0.001; \*\*\*\**P* < 0.0001. (**c**) Cellular immune response was measured using 14-colour flow cytometry panel. UMAP analysis show the (n=11) the different innate and adaptive immune cell distribution based on supervised clustering analysis. (**d**) Reduced early effector CD8^+^ T cells and NK cells after first dose of PB COVID-19 vaccination.

The second cohort (validation cohort, n=82) received 2 doses of either rAdVV or mRNA or both rAdVV/mRNA vaccines (AZ, JJ, PB, MD, or CV). This cohort was used for antibodies/cytokine measurements at large scale as well as immunophenotyping. In this cohort, samples were collected 3-12 months after two doses of vaccination from 5 different COVID-19 vaccines. Individuals were vaccinated with PB (n=34), MD (n=8), CV (n=3), JJ (n=2) and a combination of AZ and PB/MD (n=34). Further, most of the individuals had taken 3^rd^ dose either PB (n=49) or MD (n=11) vaccine to avoid any thrombotic thrombocytopenia related concerns related to the AZ vaccine^32^. Only six participants were available from CV vaccination after 3-6M PoV2 [3 participants volunteered for CureVac clinical trial (2 doses regime) before they have taken rAdVV or mRNA vaccines]. In our cohort, 3^rd^ dose of vaccinated individuals (post-vaccinated 3; PoV3) only had either PB or MD mRNA vaccines (Suppl. Table 1; Patient demographics). No prior male or female selection was performed for the recruitments. The cohort contained roughly two times female (n=74) compared to male (n=36) volunteers. The overall mean age of PoV2 (7-10 days post-vaccination 2) was 41.5 years, whilst the mean age of PoV2 (3-6M post-vaccination 2) was 37.7 years.

### Humoral and cellular immunity after first dose of vaccination (PoV1)

Previous studies have shown that by 21 days post vaccination 1 (PoV1), there is an increase in spike S1 antibody levels compared to unvaccinated individuals^33,34^. To characterize the cellular and humoral responses, we collected the blood samples (PBMCs and plasma) at 7-10 days post vaccination 1 (PoV1) and performed a 3-plex assay to identify the spike S1, Nucleocapsid and Receptor Binding Domain (RBD) IgG specific antibodies to SARS-CoV-2 virus (Suppl. Fig. 1a-c). We found that humoral response (Spike S1 antibodies levels) did not change significantly compared to pre-vaccinated individuals (PrV1) in mRNA PB vaccination (Fig. 1b). This was expected as about 14 days are generally required to develop an IgG antibody response PoV1. However, cellular immunity was impacted even at day 7-10 in the PB 1^st^ post-vaccinated (PoV1) individuals (n=7) based on 14-colour immunophenotyping flow cytometry data. We performed a uniform manifold approximation and projection analysis (UMAP) of the PrV1 and PoV1 PBMCs samples acquired on flow cytometry. We identified 8 major cell clusters including monocytes, CD4^+^ T, CD8^+^ T, NK, NKT, B cells and CD4^+^ FOXP3^+^ regulatory T cells (Tregs) based on unsupervised clustering (Fig. 1c). Supervised annotation further confirms the specific cell clusters. Further overlay UMAP plots suggest that major visual changes appeared in CD8^+^ T cells, CD4^+^ T cells, and NK cells compartments (Fig. 1d; left panel). Further supervised clustering of CD8^+^ T cells identified a major reduction in early and intermediate effectors cells (Fig. 1d middle and right UMAP panels). We observed a significant decrease (p=0.035; Wilcoxon rank-sum test) in early effector CD8^+^ T cells after first dose of PB vaccination (PoV1) and there was a trend of increased Tregs, and CD56^+^ NK cells in the PoV1 group compared with PrV1 (n=6) (Supp. Fig. 1d). No significant changes were observed for other cell types. Thus, immune subsets were affected in the first 7-10 days after the 1^st^ dose of PB vaccination.

### Altered humoral and cellular profiles after the second COVID-19 vaccination dose (PoV2)

#### I. Humoral immune response

Previous multiple studies showed that PoV2 causes an incremental increase in antibody levels after mRNA or rAdVV vaccinations, respectively^1,2,7,9,11,14,33,35^. We found a significant increase in S1 antibody levels in PoV2 (5-250 ug/ml) after the MD and PB vaccination prior vaccination (PrV1) (Fig. 1b). However, in case of AZ, JJ, and CV, it was difficult to identify the increase in antibody levels due to limited number of individuals prior to vaccination (Fig.1b). High antibody levels (5-500ug/ml) were evident between 3-6 months after PoV2 in vaccinated individuals, irrespective of vaccination platform (AZ, PB, and MD). Previous large population-based studies showed that female had higher levels of SARS-CoV-2 antibody compared with male after COVID-19 vaccination and furthermore with ageing decline in antibody levels^36,37^. Female volunteers, irrespective of vaccination platform, showed increased levels of S1 antibodies levels compared to males, although the difference was not significant (Suppl. Fig. 2a). We compared the S1 antibodies levels in different age groups and found that younger vaccinated individuals (18-29 age group) had a significantly increased (p=0.006; Kruskal-Wallis test and correct for multiple comparisons Dunn’s post-hoc test) response compared with middle aged individuals (age group 30-39; p=0.045, age group 40-49; p=0.011, age group 50-59; p=0.007), whilst older individuals (group 60+) had reduced S1 antibodies levels although this did not reach significance (Suppl. Fig. 2b). RBD antibody levels were also significantly higher in younger vaccinated individuals (18-29 age group) compared with middle age (age group 40-49; p=0.02, age group 50-59; p=0.002) (Suppl. Fig. 2c). Antibody responses were significantly enhanced after 2^nd^ dose across vaccine platforms when compared to PrV1 or first dose of vaccination and this humoral immune response was present up to 6 months, then declining from 6-8 month (PrV3) it started to decline significantly [p=0.0001; Kruskal-Wallis test and correct for multiple comparisons Dunn’s post-hoc test; p=0.0001; PoV2 (3-6M) vs PrV3 (before taking 3^rd^ dose of vaccine)] (Fig. 1b).

#### II. scRNA-seq immunophenotyping-cell classification at cellular and transcriptomics level

Using a multimodal single cell approach we further characterized the immunity after the 2^nd^ vaccine dose. Blood samples (PBMCs and plasma) from patients in the discovery cohort were collected prior to vaccination (PrV1, n=8) and 7-10 days after the 2^nd^ vaccination (PoV2) of the rAdVV vector based COVID-19 vaccines AZ (n=5) and JJ (n=3). JJ samples (n=3) were collected after the 1^st^ vaccine dose, as the approved dosage^4^ have been considered sufficient in the vaccine regimen to induce the equivalent immune response to two-dose vaccines such as rAdVV (AZ) or mRNA (BP and MD) vaccines (Fig.2a and Suppl. Table 1). Furthermore, for mRNA based COVID-19 vaccines including PB (PoV2; n=2 and PoV3; n=3), MD (n=3), and CV (n=3), we collected samples after 7-10 days of 2^nd^ dose of vaccination. We performed single-cell immune profiling including 5’ RNA sequencing, and surface protein detection with DNA-barcoded antibodies [**C**ellular **I**ndexing of **T**ranscriptomes and **E**pitopes by **Seq**uencing (CITE-seq)] on a 10x Chromium platform (Fig. 1a and Fig. 2a). After filtering of low-quality cells, a total of 329,920 cells were included in the scRNA-seq analysis (Suppl. Fig. 3, 4). In total, 27 clusters were obtained, after applying Louvain clustering to a shared nearest-neighbor graph constructed with the first 50 principal components on the normalized scRNA-seq gene expression matrix. The clusters were visualized in UMAP space (Fig. 2b). Each cluster was manually assigned to a specific cell type based on gene expression markers (RNA) and surface protein levels (protein) as listed in Suppl. Tables 2&3 (Fig 2b, c and Suppl. Fig. 5). Further, we used the Azimuth (reference-based cell annotation of PBMCs)^38^ to confirm the cell type for each of the cell clusters that were manually annotated with gene expression markers (Fig. 2b, c). The manual annotation matched the Azimuth cell annotation for most of the cell clusters (Suppl. Fig. 6). The majority of the cells (30-40% of total PBMCs) were identified as CD4^+^ T naïve (cluster 0; CD4^+^ TN), central memory (TCM) and effector memory (TEM) CD4^+^ T cells (clusters; 1, 10, 17, 15 and 22) (Fig. 2). Cluster 15 comprises regulatory T cells (Tregs) based on Tregs unique marker genes including FOXP3, IZKF2, IL-2RA, RGS1 and CTLA-4^39^. Important to note that CD4^+^ IL-7R^+^ cells were further divided into different subsets based on CCR7, SELL, AQP3, CD27, CD28 mRNA or surface protein identification (Fig. 2c, Suppl. Table 1; Cluster genes, Suppl. Fig. 6a)^40^. However, most of TCM CD4^+^ T cell clusters contain a mixture of the same characteristics genes which are difficult to specify into sub-groups cells therefore we defined as TCM I, TCM II and TCM III based on relative gene expression levels (Suppl. Fig. 6, Suppl. Table 4; cluster gene). Four CD8^+^ T cells subsets (20-25% of total PBMCs) were identified, including TN (cluster 6), TCM (cluster 8), TEM I and TEM II (clusters 4 and 5 respectively) (Fig. 2b-c and Suppl. Fig. 6b). NK cells (10-12%), B cells (6-8%) and monocytes (12-15%) makes the remaining percentage of volume of PBMCs (Fig. 2b, c). Overall, based on unsupervised clustering, we can identify the 25 major types based on differential gene expression and percentage of individual cell types. Moreover, these cell clusters percentage classification analyses expanded to distinguish the differences among pre-vaccination *vs*. post-vaccination (2^nd^ or 3^rd^ dose of vaccine) for two major vaccine groups (rAdVV and mRNA) and also for the individual vaccines (AZ, JJ, MD, CV and PB) later in discrete sections.

**Fig 2.**
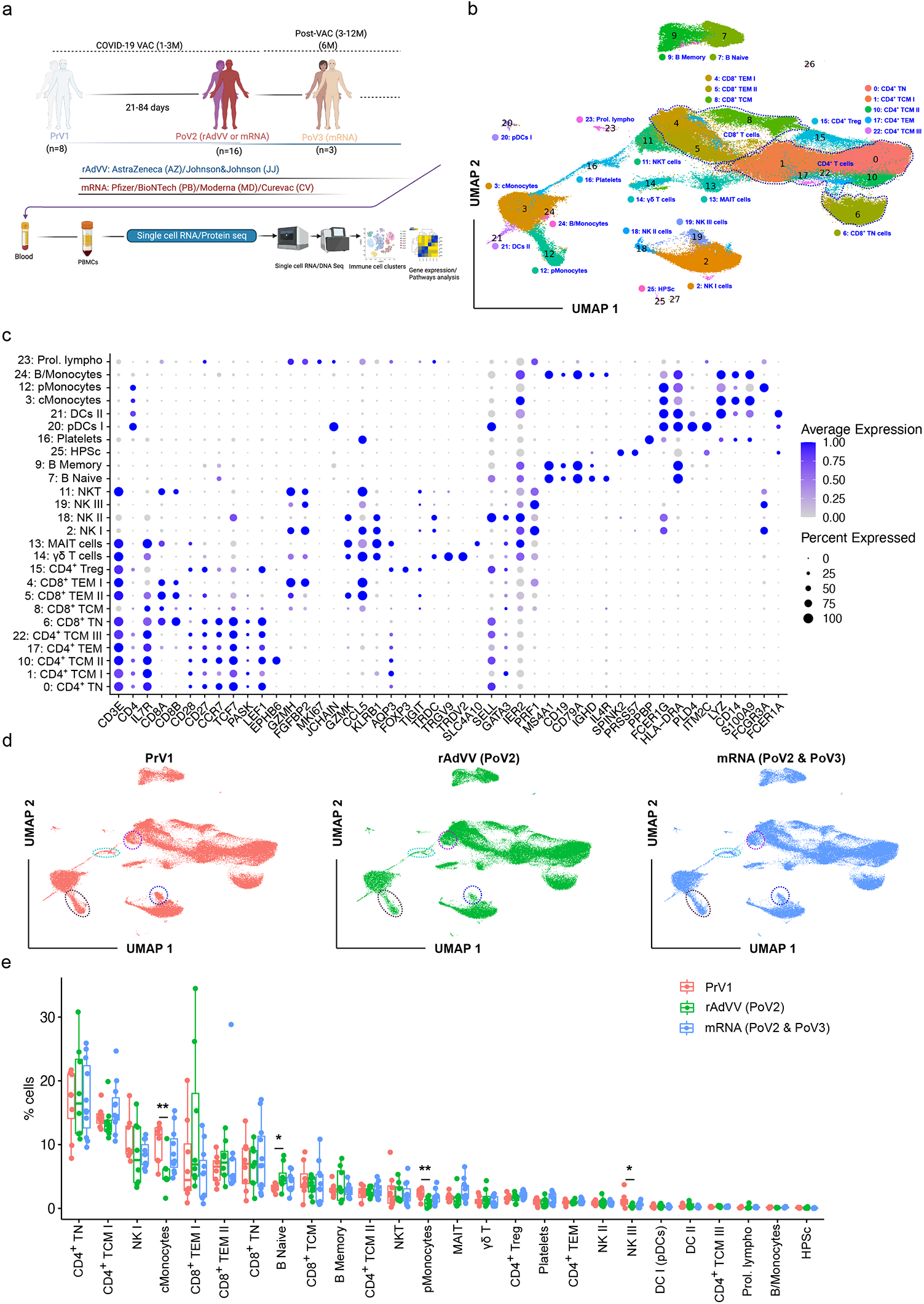
Cellular immune response after 2^nd^ dose of rAdVV and mRNA COVID-19 vaccination. **(a)** Experimental design for the multi-omics single-cell profiling. Blood was collected from naive individuals before vaccination (PrV1), 7-10 days after dose 1 (PoV1; n=8), 7-10 days after dose 2 (PoV2; n=8 rAdVV, n=8 mRNA, and 7-10 days post V3 (PoV3; n=3 PB). Longitudinal PBMC samples (PrV1, PoV2, and PoV3). scRNA-seq and antibody-based surface protein detection (130 marker proteins) was performed. **(b)** Uniform manifold approximation and projection (UMAP) of the single-cell RNA-seq profiles for all analysed samples. The Louvain algorithm was used for detecting different cell clusters from a shared nearest neighbour graph based on the first 50 principal components of the RNA expression matrix. Major subsets of CD4^+^ and CD8^+^ T cells are highlighted in dotted line circles. (**c**) Dot plots of key gene expression markers used for cluster cell type annotations. (**d**) UMAP representation of the scRNA-seq data before vaccination (PrV1), after rAdVV-based vaccination (PoV2; n=8) and mRNA-based vaccination (PoV2 and PoV3; n=8) based vaccination. An equal number of cells per group are presented for visual comparisons (n=77412 cells/group). (**e**) Box plots comparing percentage of different cell subsets across each group. Wilcoxon rank-sum test for two sample comparisons for PrV1 vs PoV2 (rAdVV and mRNA) or PoV3 for individual cell clusters. Box plots indicate medians and interquartile ranges. \**P* < 0.05; \*\**P* < 0.01; \*\*\**P* < 0.001; \*\*\*\**P* < 0.0001.

#### III. scRNA-seq immunophenotyping - comparative analysis of rAdVV and mRNA-based COVID-19 vaccines at single cell transcriptomic level after 2^*nd*^ *dose*

We performed a differential cell abundance testing between before (PrV1) and after the 2^nd^ dose of rAdVV (PoV2) vaccines as well as before and after the 2^nd^/3^rd^ dose of mRNA-based (PoV2/3) vaccines using a Wilcoxon rank-sum test (Fig. 2d-e). We found that cMonocytes (p=0.006), pMonocytes (p=0.04) and NK III (p=0.02) were significantly reduced whilst naïve B cells were significantly increased (p=0.03) after rAdVV vaccination (PoV2) (Fig. 2d, e). cMonocytes, pMonocytes and NK III cells were also reduced after mRNA vaccination (PoV2/3) whilst B naïve cells were increased (similar trend equivalent to rAdVV vaccine) although the difference was not significant for this vaccine group (Fig. 2d, e).

When considering individually the vaccine groups, we undertook a comparative global analysis among the PrV1 with individual AZ (PoV2), JJ (PoV1), MD (PoV2), CV (PoV2) and PB (PoV2/3) vaccines. We observed that cMonocytes, pMonocytes, NK III were significantly decreased whilst CD4^+^ TCM III cells were significantly increased after AZ (PoV2) vaccination based on Wilcoxon rank-sum test (Fig. 3a, b). Naïve B cells, γδ T cells and proliferating lymphocytes were significantly increased after JJ (PoV1) vaccination (Fig. 3a, c). After MD (PoV2) vaccination, CD8^+^ naïve T cells and platelets were significantly increased, whilst the CD8^+^ TEM I cell population was decreased (Fig. 3a, d). Proliferating lymphocytes were only increased after CV (PoV2) vaccination (Fig. 3a, e). After PB (PoV2/3) vaccination, no significant changes were observed, but a trend of increased CD4^+^ TCM I and decreased NK cells (Fig. 3a, f). Tregs tended to increase after mRNA COVID-19 vaccination (Fig. 3g, Suppl. Fig. 7). Interestingly, platelets tended to decrease in rAdVV (PoV2) vaccines whilst tended to increase in mRNA (PoV2/3) (Suppl. Fig. 7). Taken together, our single-cell analysis after vaccination pointed out that monocytes, T and NK cells could be an important marker in vaccine induced signature detection.

**Fig. 3.**
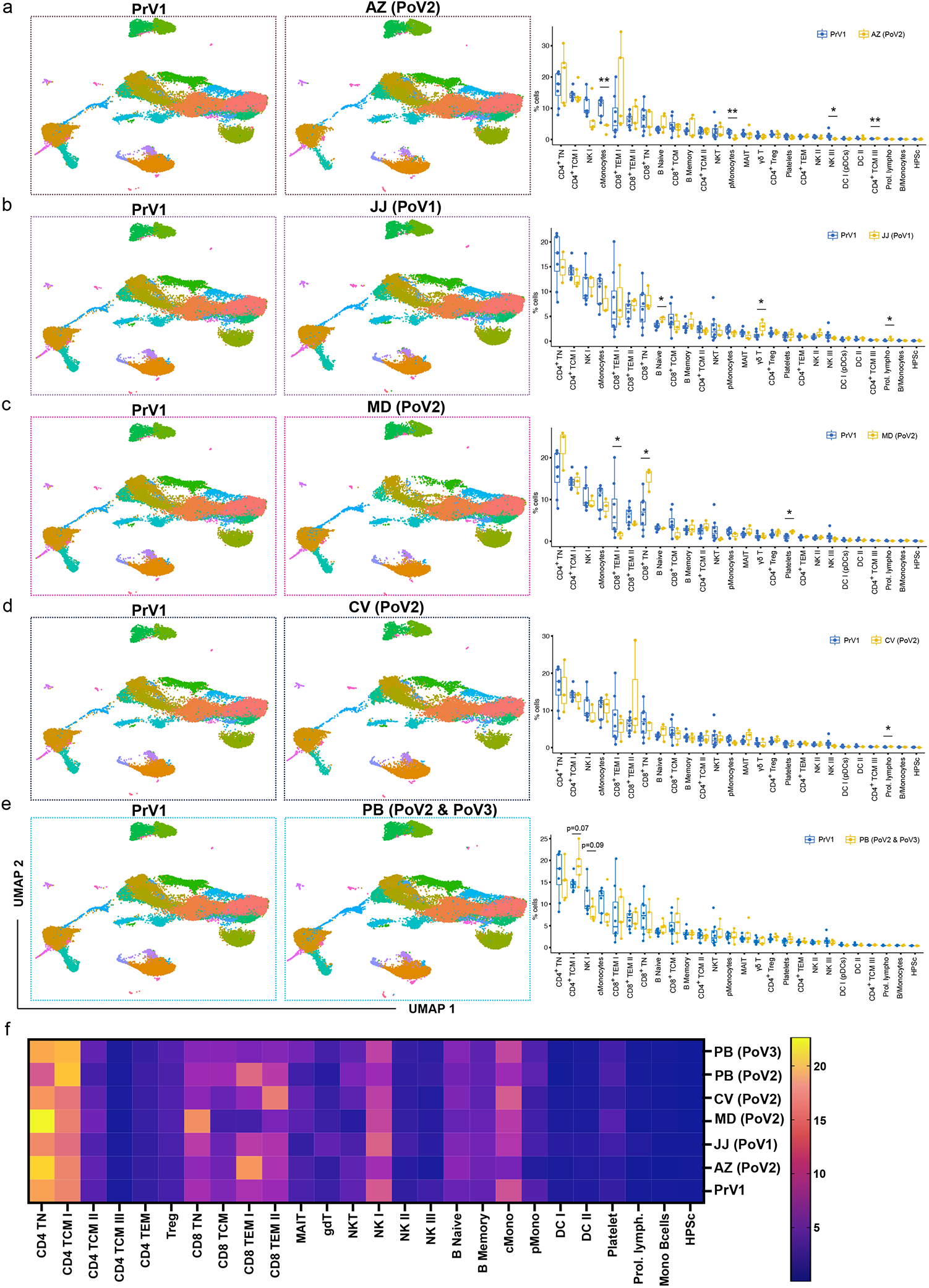
Cellular immune response after 2^nd^ or 3^rd^ doses of individual COVID-19 vaccines. (**a**) UMAP cell clusters in PrV1 (left; n=8; cells=85710) and PoV2 (middle; n=5; cells=43553) in AZ vaccination. Box plots represent changes in different immune cell subsets post-vaccination (right panel). (**b**) PoV1/2 (middle; n=3; cells=33859) in JJ vaccination. Box plots represent the percentage of the immune cell subsets pre- and post-vaccination. (**c**) PoV2 in MD (middle; n=3; cells=36493) vaccination and box plots show change in different immune cell subsets post-vaccination. (**d**) PoV2 in CV (middle; n=8; cells=24466) vaccination and box plots represent change in different immune cell subsets post-vaccination (lower panel). (**e**) PoV2 in PB (middle; n=5; cells=48250) vaccination and bar diagrams represent change in different immune cell subsets post-vaccination. (**f**) Heatmap shows the cell type percentage for each cell subset and individual vaccines groups. Wilcoxon rank-sum test for two sample comparisons (a-e) for PrV1 vs PoV2/PoV3 (rAdVV and mRNA) for individual cell clusters. Box plots denote medians and interquartile ranges. \**P* < 0.05; \*\**P* < 0.01; \*\*\**P* < 0.001; \*\*\*\**P* < 0.0001.

#### IV. Altered gene expression and signalling pathways after rAdVV or mRNA COVID-19 vaccination

We then investigated the differential gene expression before (PrV1) and after vaccination (PoV2) in pMonocytes, platelets, NK cells (type III) and NKT cell types (Suppl. Fig. 8). Most importantly majority of genes in pMonocytes for both the rAdVV and mRNA vaccines were downregulated (Suppl. Fig. 8a). To understand further the role of these genes in biological functions, we employed Metascape tool^41^ for interpretation of systems-level studies for the inference of enriched biological pathways and protein complexes contained within OMICS datasets. Pathway and gene ontology (GO) analyses derived from Metascape of downregulated genes (upregulated DEGs were not enough to have any significant change in any pathway) in pMonocytes after rAdVV vaccination revealed that they were genes involved in the regulation of CD4-positive, alpha-beta T cell activation, leukocyte chemotaxis and IL-18 signalling. Whilst in mRNA-based vaccination, downregulated genes in pMonocytes after vaccination were found to be involved in proinflammation, profibrotic mediators and NFkB signalling pathways (Suppl. Fig. 8a; right hand side figure). Several genes such as HLA-DQB1 (MHC II), CD86 (co-stimulation), CD83 (co-stimulation), CXCR4 (chemokine attracting CD4^+^ T cells) were downregulated after rAdVV vaccines compared with PrV1 whilst they were not differentially expressed after mRNA vaccination. These genes play a pivotal role in the immune system by presenting peptides derived from extracellular protein, chemoattraction and co-stimulation of CD4^+^ T cells, therefore, downregulation of these genes could be implicated in poor antigen presentation by macrophages after rAdVV vaccine. Thus, it appears that two distinct vaccine platforms (rAdVV and mRNA vaccines) have divergent pathways to regulate the CD4^+^ T cells stimulation after vaccination.

In NK III cells, pathway analysis based upon upregulated genes highlighted that NK cells could have a capacity of higher leukocyte mediated cytotoxicity, regulation of calcium ion, cellular response to cytokines, intracellular signal transduction and actin dynamics after rAdVV (PoV2) vaccination in NK cells. Whilst we could not get any enriched pathways for mRNA (PoV2) vaccine due to limited differentially expressed genes. We inferred that after rAdVV vaccination could have heightened NK cell function whilst NK cell functions are not changed with mRNA vaccination (Suppl. Fig. 8b).

In NKT cells, almost equal number of upregulated genes after both the vaccine platforms compared with PrV1 state. Pathways analysis for the common gene signature in both, the rAdVV and mRNA vaccinated cohort, revealed that both the vaccines induced estrogenic signalling, integrin mediated cell adhesion, MAPK signalling cascade, and regulation of lymphocytes activation in NKT cells. Natural killer or innate lymphoid cells inherently express more estrogenic genes^42^ (FOS, GNAI3, GNAS, JUN, KRT10, ZFP36L2, DUSP1, CYBA, ATP5F1D, CD8, DUSP1, DUSP2, CAPNS1, KLF2, RPS2 and TAF10), however, both the vaccines able to enhance this signalling pathways gene showed common non-specific modulators (cytokines or chemokines) could control the activation NKT cells (Suppl. Fig. 8c). Further, pathway analysis based on unique upregulated genes only for rAdVV vaccine suggested that these genes could be involved in heightened adaptive immune response, nuclear receptors meta-pathways, and signalling by Receptor Tyrosine kinases in NKT cells. Furthermore, T cell receptor (TCR) genes such as TRAV8-4, TRAV26-2, TRBV28, TRBV30 and TRBV4-2 were also upregulated after rAdVV vaccination. TRAV8-4 genes found be expanded in COVID-19 convalescent^43^, thus it could be reflecting that rAdVV vaccination have spike specific NKT cell activation. In case of mRNA (PoV2/3) vaccines based upon upregulated genes, pathway analysis revealed that NKT cells should be positively regulating activation of other cell types including platelets activation and aggregation, cell adhesion, cellular defence response, chemokine signalling pathways, and infection of human immunodeficiency virus 1. Most importantly, PRF1, TBX21, CD69, CX3CR1, KLRG1 genes were abundant in mRNA vaccination compared with rAdVV vaccines. These results suggested that NKT cells should have more inflammatory, activated, and cytotoxic functions after mRNA vaccination. In addition to this, TRAV6-5 gene was also upregulated in mRNA vaccines so implied the spike specific NKT cell response. Thus, differential gene expression analysis suggested that each vaccine type (rAdVV or mRNA) in individual cell cluster could have different immune response outcome, thus overall immune response should be different depending upon COVID-19 vaccination.

Dysregulated functions of platelets were involved in the vaccine-induced immune thrombotic thrombocytopenia (VITT)^44^. In platelets, rAdVV and RNA vaccination, *FOS, JUN*, and *DUSP1* were commonly upregulated. Platelets from rAdVV vaccinated individual appears to have increased inflammatory and cytotoxic phenotype based on *IER2, TNF, PRF1, GZMH*, and *EGR1* gene expression, whereas these genes were not differentially changed after mRNA vaccination (Suppl. Table 5). Therefore, we can speculate that platelets are activated after rAdVV vaccination whilst mRNA vaccines do not interfere/control their normal functions. Based on our results, it would be certainly interesting to understand the molecular functions whether platelets were hyperactivated which led to VITT after rAdVV vaccine. Furthermore, Gene pathway enrichment analysis revealed that these genes could also be involved in the regulation of CD8 TCR downstream signalling pathways by modulating MHC I expression as described previously^45^.

#### V. Dysregulated gene expression and signalling pathways after AZ (rAdVV) COVID-19 vaccination

Based on these different gene expression patterns after rAdVV or mRNA based COVID-19 vaccination, we focussed further on the individual COVID-19 vaccine to understand how differentially expressed genes could reveal the regulation of different cellular pathways in each cell cluster population which could be paramount for fundamental understanding in immune response and vaccines. Thus, we focussed on AZ vaccine as we had most significant cell types were changed in PrV1 vs AZ (PoV2).

Comparing pMonocytes in PrV1 individuals with AZ (PoV2), we found that 188 genes were differentially regulated which allowed further gene enrichment analysis (Suppl. Table 6). Gene enrichment pathway analysis in pMonocytes pointed out modulation of the oxidative phosphorylation (metabolic function), cardiac muscle contraction and mitochondrial biogenesis pathways were upregulated in AZ vaccinated individuals. Whilst, based on downregulated genes in pMonocytes, we observed that NFkB signalling, IL-18 signalling, T cell activation (AHR, ANXA1, CD86, CDKN1A, CTNNB1, CD55, HLA-DQB1, IL1B, LYN, SMAD7, PNP, PLEK, CCL3, TNFAIP3, NR4A3, RIPK2, CD83, PELI1, ZMIZ1, SELENOK, SAMSN1, NFKBIZ, NLRP3, TICAM1), TCR signalling (HLA-DQB1, NFKB1, NFKBIA, UBC, RIPK2, LCP2, GNA13, TNFAIP3, IL1B, VIM, CD83), wound healing, IL-10 signalling, and VEGFA-VEGFR2 pathways were downregulated (Suppl. Fig 9a). We also examined the common gene expression in other cell types including CD4^+^ TCM III, cMonocytes, γδ T cells and Tregs which could be important for shaping the immune response and eliciting effector immune (Suppl. Fig. 9).

In comparative analysis among different immune cell subsets, we interestingly found that *GADD45B* and *IER2* genes were upregulated after AZ (PoV2) vaccine compared with PrV1 individuals, whilst in mRNA (PoV2) vaccination (MD and CV) no differences were discerned for the same genes. Stress response gene GADD45B appeared to be induced by environmental stress or treatment with DNA-damaging agents and involved in DNA methylation thus control adult neuronal differentiation^46^. Thus, we speculate that epigenetic changes may occur due to upregulated expression of GADD45B after rAdVV vaccines as described earlier for other infectious disease vaccines^47^. Additionally, there were some common unique gene signatures in monocytes cell subsets such as *IL1B, CD83* and *IER3*. These three genes were downregulated after AZ, CV and MD vaccination compared with PrV1. Overall, our single cell multi-OMICS analysis revealed rAdVV and mRNA vaccines have different mechanism of action for activation of lymphocytes and monocytes, respectively.

### Altered humoral and cellular profiles across different COVID-19 vaccines post-vaccination 3 (PoV3)

Furthermore, third dose of PB vaccine mRNA vaccination was given in previously fully vaccinated individuals (PoV2) after 6 months with AZ or PB. We noticed again improved humoral response (spike S1 antibody levels) in PoV3 vaccinated individuals compared with PoV2 vaccinate individuals (Fig. 1b). Further cellular response was verified using scRNA-seq and multi-colour flow cytometry panel staining. Surprisingly, based on comparison among PrV1 and PoV3, we found that Tregs were tended to be increased in PoV3 vaccinated individuals (p=0.08; Wilcoxon rank-sum test) (Suppl. Fig. 7c). Furthermore, total CD8^+^ T tended to decrease and increase of CD4^+^ T cells when comparisons were made among PrV1 and PoV3 (Fig. 2f and Fig. 3g).

### Antigen specific T cell response is affected with rAdVV or mRNA-based vaccines

The effect of vaccine immunization was first explored on recipient T cell populations in a cohort of 28 healthy volunteers who had not been previously infected with SARS-CoV-2 (discovery cohort), and a validation cohort (n=82) after 3-6 months of vaccination, as well as 3^rd^ or 4^th^ booster dose of mRNA vaccines. By collecting longitudinal peripheral blood samples before immunization, one week following booster immunization (day 7), and at an additional time point approximately 3 weeks to three months later (day 27 - 84), we captured signatures of the initial immune response to PB mRNA vaccines as well as lasting immunity. To identify the antigen-specific T cell stimulation (either CD4^+^ or CD8^+^ T cells), we used 1×10^6^ PBMCs PrV1, PoV2 or PoV3, cultured them for overnight in presence of spike peptide, and performed cytokine assay analyses (Fig 4a). After PB vaccination, an increase of spike-specific T cells was present (Fig. 4b). MD vaccine caused similar results, however, AZ vaccinated individuals had rather low expression of spike specific T cells (Fig. 4b). Data from JJ were not completely conclusive due to a limited number of samples; one individual had already activated expression of spike specific T cell response, which was decreased after vaccination, whilst in other individuals we could not detect enough amount of spike-specific T cells (Fig. 4c).

**Fig. 4.**
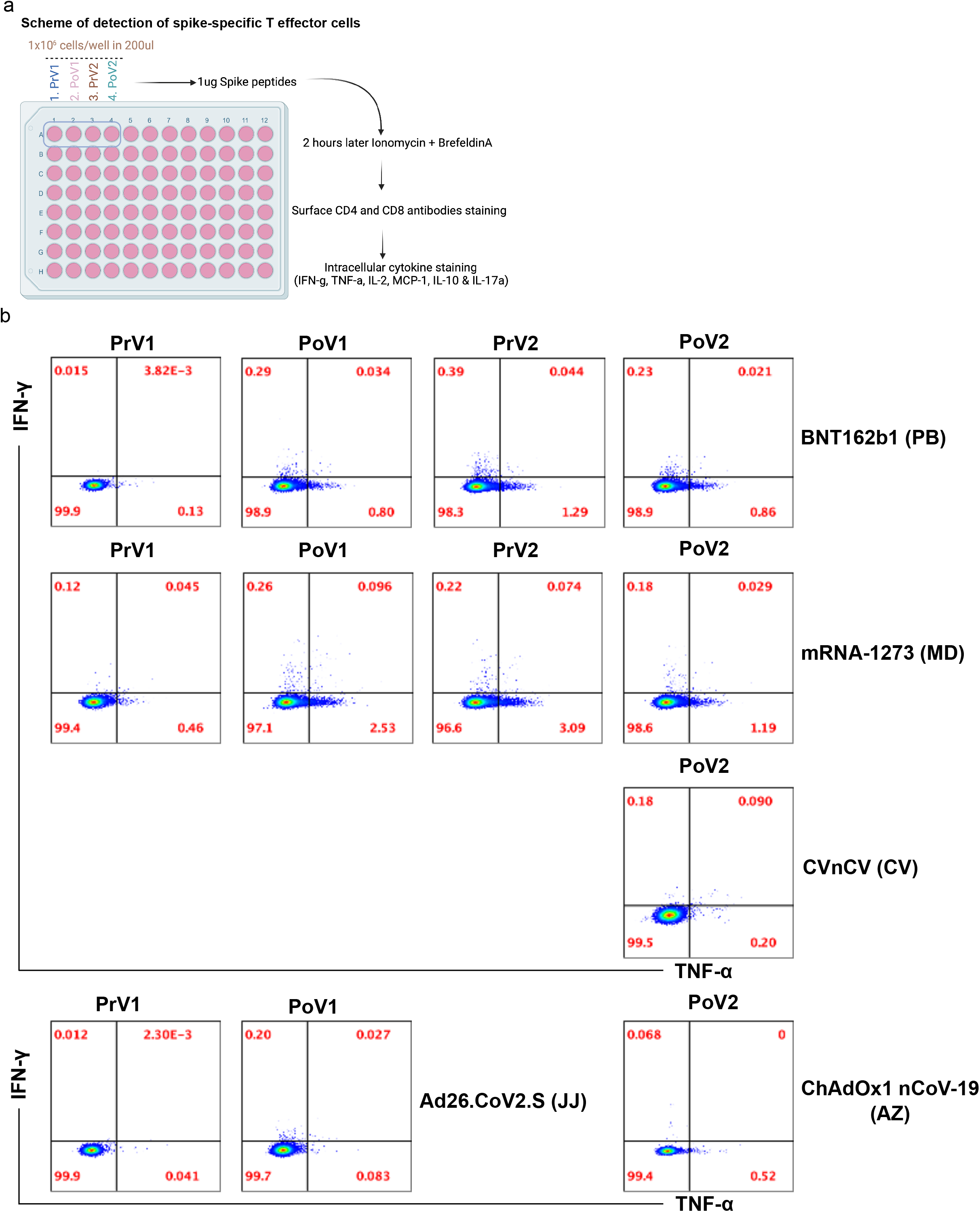
Detection of antigen-specific (spike-antigen) T cells response in PoV2 vaccinated individuals with SARS-CoV-2 mRNA or rAdVV vaccination. (**a**) A cartoon is presented the experimental strategy for detecting antigen-specific T (CD4^+^ T cells) (**b**) Intracellular flow cytometry was performed for IFN-γ and TNF-α after treatment with 1ug/ml of peptide for overnight. PoV1 and PoV2 for MD and PB show strong IFN-γ and TNF-α positive CD4^+^ T cells, whilst, CV had a few IFN-γ^+^ or TNF-α^+^ CD4^+^ T cells after 2^nd^ dose of vaccination. (**c**) IFN-γ and TNF-α staining after was 1ug/ml of peptide treatment for overnight for rAdVV COVID-19 vaccines. PoV1 and PoV2 for JJ and AZ show a few IFN-γ^+^ or TNF-α^+^ CD4^+^ T cells. (**d**) Summary statistics of all the experiments to determine the antigen-specific T cell response. Bar diagram change in different immune cell subsets post-vaccination (lower panel). Wilcoxon rank-sum test for two sample comparisons or multiple samples comparison was performed using Kruskal Wallis test and post-hoc Dunn’s test for multiple comparisons for PrV1 vs PoV2 (rVV and mRNA) or PoV3 for individual cell clusters. Error bars denote medians and interquartile ranges. \**P* < 0.05; \*\**P* < 0.01; \*\*\**P* < 0.001; \*\*\*\**P* < 0.0001.

#### Altered signalling cascades in T cell activation and their role in antigen presentation

To validate further whether different vaccines may induce differential T cell antigen presentation, we mined our scRNA-seq data set for the genes relevant for T cell signalling and antigen presentation. First, we investigated the expression of CD48, CD2, and CD244 genes which are responsible T cell antigen presentation and effector functions^48–51^. CD244:CD48 interactions regulate target cell lysis by NK cells and CTLs, which are important for viral clearance and regulation of effector/memory T cell generation and survival^50,52^. CD48 is involved in T cell and antigen presentation and ligand for CD48 is either CD2 or CD244^48,49,51^. We opted AZ and PB vaccines for a direct comparison due to limited number of samples in other groups. We noticed decreased expression of *CD2, CD58, CD244* and *CCR5* at mRNA and protein level whilst increased expression of *CD48* and long non-coding RNA (lncRNA) *MALAT1* in vaccinated individuals with AZ compared with PrV1 as well as with PB (Fig. 5a & b and Suppl. Fig. 10). Furthermore, *ITGB2* (CD18) protein is involved in cell adhesion, cell surface signalling and immune (CD11a/CD18) together ICAM-1 (CD54) synapse formation for antigen presentation^53^. We found that ITGB2 (CD18) was upregulated in pMonocytes (Suppl. Fig. 10), which could be responsible for governing the antigen presentation and outcome of cytotoxic function of immune cells. Further, when we compared the expression of *CD2, CD244* and other cell activation genes at a global level, we yielded the similar outcomes as well for rAdVV and mRNA COVID-19 vaccine compared with PrV1. However, MALAT1 expression was slightly reduced in CD8^+^ TCM, MAIT and NK III cells in rAdVV and mRNA vaccinated individuals (Suppl. Fig. 11). Thus, our data suggest that different mechanisms are operating for activation of immune cells by two different types of COVID-19 vaccination.

**Fig. 5.**
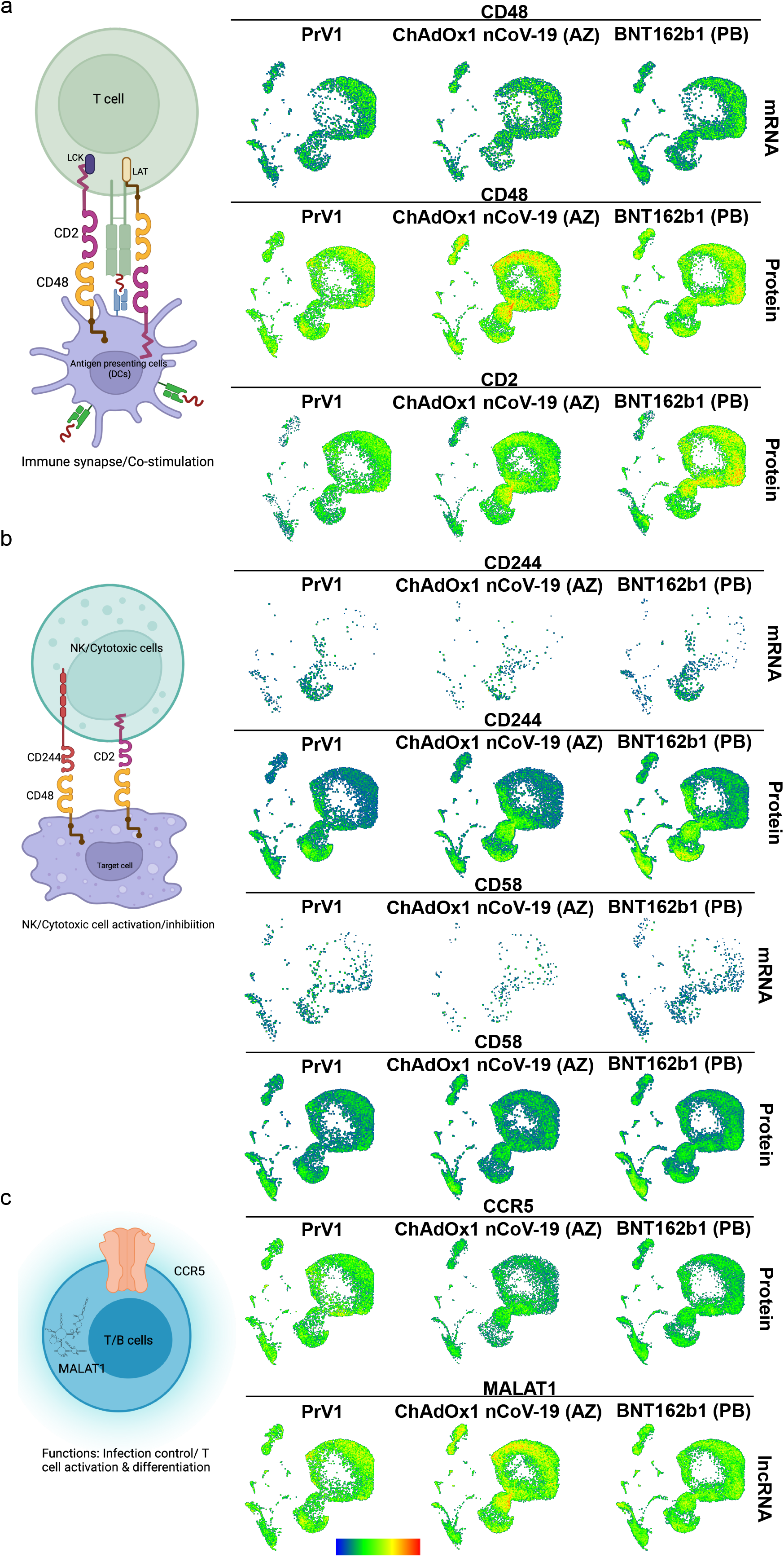
2^nd^ dose of AZ (rAdVV) vaccination induces a distinct immune signature compared with PrV1 and PB vaccines through CD48-CD244-CD2-CD58-MALAT1. (**a**) Global gene and protein expression of CD244, CD48, CD2 and CD58 on all the immune cell subsets. (**b**) RNA and protein expression of CCR5 and RNA levels of lncRNA – MALAT1 in PBMCs cell subsets.

### Vaccination (2^nd^ and/or 3^rd^ dose) increased the percentage of CD4^+^ TCM cells and reduced NKT cells

CD4^+^ T cells play an important role in helping B cells for the generation of antibody producing B cells. First, we pooled samples of lymphocytes for mRNA vaccination of different time points and compared the immune dynamics after 2^nd^ and 3^rd^ dose (both before and after vaccination). We performed the UMAP analysis to identify the cell distribution of T, B, and NKT cells and of cell specific markers (Fig. 6a & Suppl. Fig. 12). We observed a clear difference in CD4^+^ T cell and NKT cell distribution after 2^nd^, 3^rd^, and 4^th^ dose of mRNA vaccination (Fig. 5b & c). Further, 2D analysis suggested that overall CD4^+^ T cells were increased significantly after 3^rd^ dose and CD4^+^ TCM memory cells were significantly decreased (Fig. 6d-e). CD8^+^ T cells tend to increase after the 4^th^ dose, and TEMRA cells appeared to be decreased. TCM CD8^+^ T cells tend to increase after PrV1 to PoV2, but no increase was detected after 3^rd^ or 4^th^ dose, respectively. Furthermore, CD8^+^ TEM tended to be decreased (Fig. 6d). More interestingly, NKT cells were significantly different after 3^rd^ vaccination when compared with PrV3 individuals (Fig. 6f). Overall, one can conclude that vaccination most significantly changes the NKT and T cell compartments as well.

**Fig. 6.**
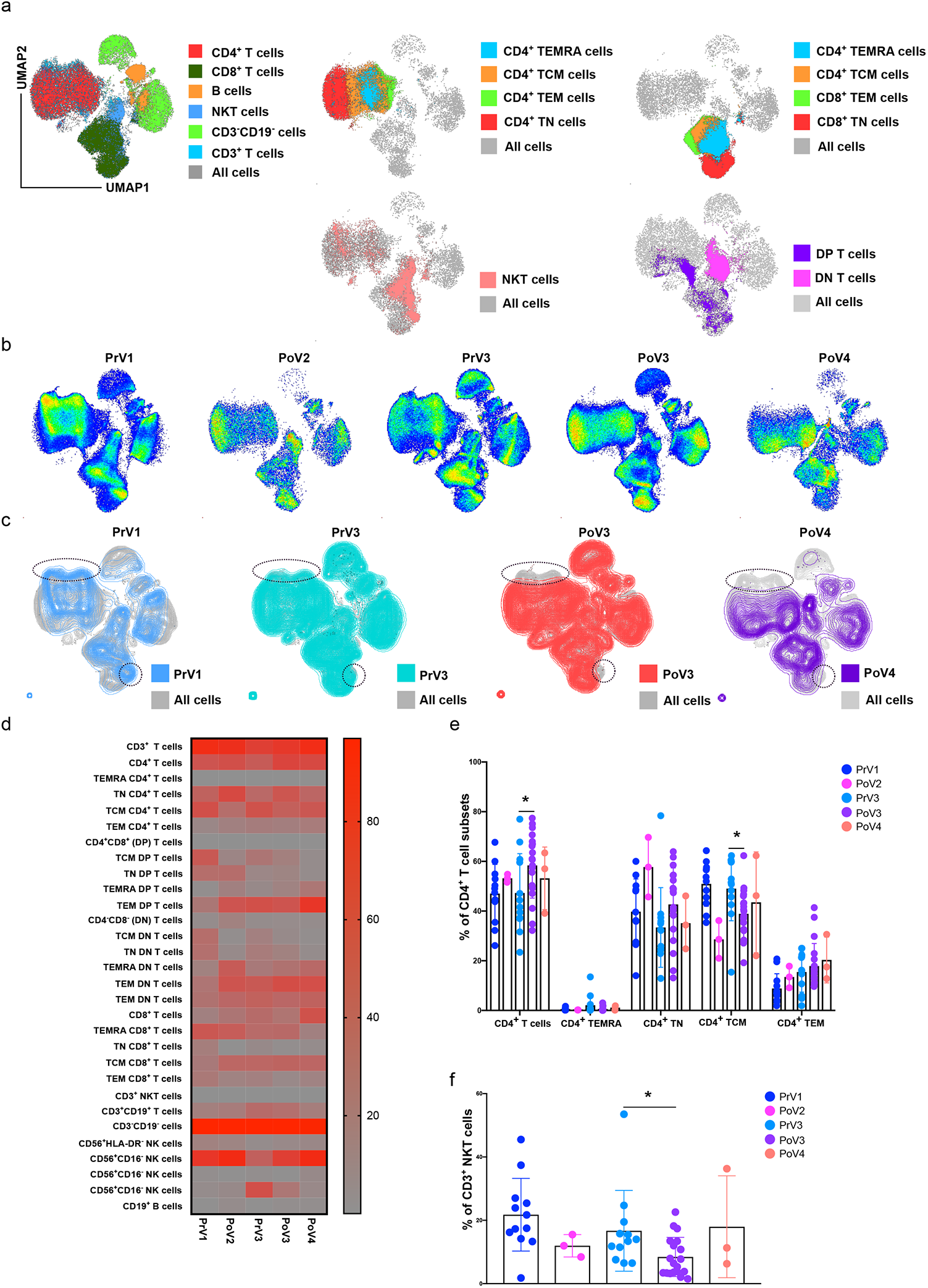
Increase in CD4^+^ T cells and reduced CD4^+^ TCM and CD3^+^ NKT cells PoV3 vaccination. (**a**) Total 44 samples were concatenated for lymphocytes and used for unsupervised clustering analysis using UMAP dimension reduction method. Cells were clustered into major cell subsets including CD4^+^ T cells, CD8^+^ T cells, CD4^+^ NKT cells and CD8^+^ NKT cells, B cells and CD3^-^CD19^-^ (mostly NK cells). Each cell population was colour coded as shown in fig (left figure). Various CD4^+^ T cells, CD8^+^ T cells and NKT cell subsets were depicted (right figure, 4 different UMAP plots). (**b**) Individual UMAP plots for Pre-VAC1, PoV2, Pre-VAC3, PoV3 and PoV4 vaccinated individuals. (**c**) Overlay of individual vaccinated group compared with all cells. Major visual difference was detected in CD4^+^ T cells and NKT cell population. (**d**) Heatmap plots to summarize all the data to determine the role of individual lymphocyte subsets. (**e**) Bar plots from CD4^+^ T cell subsets and major statistical difference was reached for total CD4^+^ T cells as well as CD4^+^ TCM in PrV3 and PoV3. (**f**) Bar plots from CD4^+^ T cell subsets and major statistical difference was reached for total CD3^+^ NKT cells in PrV3 and PoV3. Wilcoxon rank-sum test for two sample comparisons or multiple samples comparison was performed using Kruskal Wallis test and post-hoc Dunn’s test for multiple comparisons for PrV1 vs PoV2 (rAdVV and mRNA) or PoV3 for individual cell clusters. Error bars denote medians and interquartile ranges. \**P* < 0.05; \*\**P* < 0.01; \*\*\**P* < 0.001; \*\*\*\**P* < 0.0001.

### Cytokines could be a biomarker signature in post-vaccinated (PoV2: 3-6M) individuals

Cytokines (e.g. IL6) and chemokines (e.g. CXCL-10) are involved in B cells or plasmablast activation, and age could a determining factor for cytokine production^54,55^. Further, a recent study pinpoints that CXCL-10 levels were upregulated after first dose of mRNA (PB) vaccination^56^. Applying a 13-plex cytokine assay, we identified that MCP-1, IL-10RA, and CXCL-10 were significant increase in the post-vaccinated cohort (3-6M) (Fig 7a). Therefore, we compared the data of PrV1 vs PoV2 (3-6M) groups and found that MCP-1, IL-1RA, and CXCL-10 were significant increased (Fig 7a). Further, comparing data with all different vaccinations we did not find any specific vaccine-related changes but rather global changes (Suppl. Fig. 13). Age-based analyses suggested that older individuals (49^+^ age and 60^+^ age groups) had reduced levels of CXCL-10 compared with 18^+^ age group individuals (Fig. 7b). Further, we attempted to identify the correlation among cytokines levels and vaccine-induced antibodies. Our antibody and cytokine correlation analysis revealed that IL-6, IFN-α2, IL-2, IL-7, CCL3, and CXCL-10 were mildly positively correlated, whilst IL-10, CXCL-8, TNF-α, IFN-γ, and G-CSF have a weak negative correlation with Spike S1 antibody levels (Fig. 7c). RBD antibody levels were negatively correlated with MCP-1 cytokine levels (Fig. 7c). Overall, our data revealed that MCP1, IL-10RA and CXCL-10 cytokines could potentially be the vital cytokines which can be used for detection of vaccine efficacy.

**Fig. 7.**
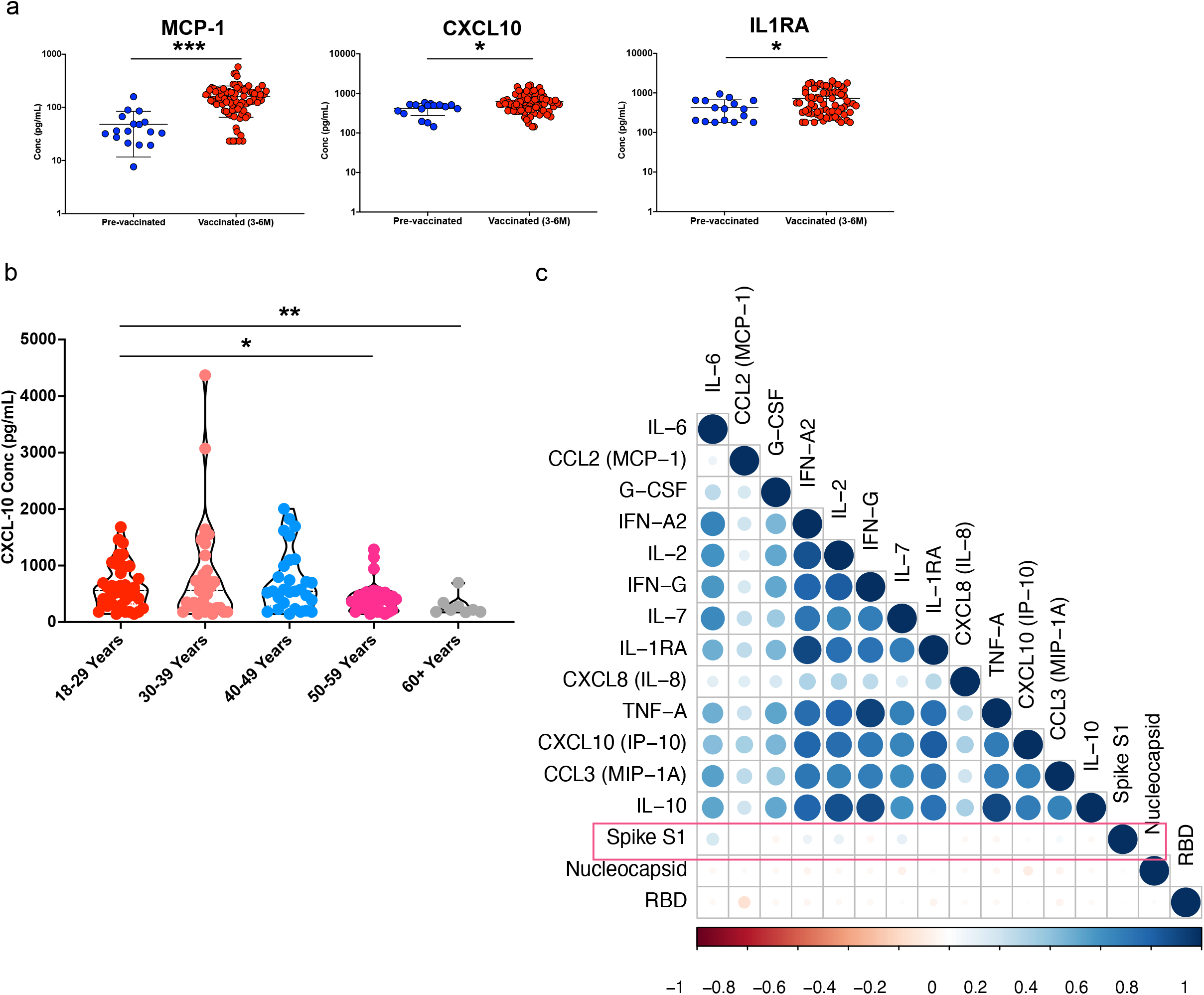
Increased MCP1, IL1RA and CXCL-10 PoV2 in 3-6 months vaccinated individuals. (**a**) 13-plex flowcytomix assay was used for detection of humoral immune response and 3 cytokines (MCP1, IL1RA and CXCL10) were significantly upregulated compared to PrV1 samples. Wilcoxon rank-sum test for two sample comparisons was used for data analysis [PrV1 vs PoV2 (3-6M)]. Error bars denote medians and interquartile ranges. \**P* < 0.05; \*\**P* < 0.01; \*\*\**P* < 0.001; \*\*\*\**P* < 0.0001. (**b**) CXCL-10 levels among different age vaccinated individuals. Wilcoxon rank-sum test for two sample comparisons was used for data analysis. Age group 18-29 used as baseline for comparing with other age groups Error bars denote medians and interquartile ranges. \**P* < 0.05; \*\**P* < 0.01; \*\*\**P* < 0.001; \*\*\*\**P* < 0.0001. (**c**) Pearson correlation analysis among cytokines and antibodies levels.

## Discussion

Single cell-based analysis tools offer new ways to compare how individual patient’s cells respond to treatment or change during infection. The method could help to monitor immune response after vaccination or against infections. Here, we demonstrate the comparative analysis of 5 individual vaccines against SARS-CoV-2 [ChAdOx1 nCoV-19; AstraZeneca (AZ), Ad26.CoV2.S; Johnson & Johnson (JJ), BNT162b1; Pfizer/BioNTech (PB), mRNA-1273; Moderna (MD), CVnCV; CureVac (CV)] which can be divided broadly into two different groups (rAdVV and mRNA based vaccines) at single cell RNA, proteogenomic, and protein levels. We describe that rAdVV (AZ) and mRNA (PB) vaccines work differently and activate immune cells contrarily at gene levels. Additionally, our study highlighted that innate and adaptive cell such as monocytes, NKT, B and T cells play an orchestrated role in the generation of effector immune response as demonstrated after different doses of vaccination compared with PrV1.

Major observations are: (1) No change in humoral response of 1^st^ dose of COVID-19 vaccination was observed for any vaccination after 7 days, however a decrease in effector CD8^+^ T cells and increased NK cells (mRNA PB) were observed. (2) After a 2^nd^ dose of COVID-19 vaccination (7-10 days) a strong humoral immune response irrespective of vaccination (AZ, PB and PB), but only a weak response after JJ (PoV1 but considered equivalent to 2 doses). Cellular immune response was robust which led to overall increase in Tregs (mRNA-PoV3). Decrease in monocytes (pMonoyctes and cMonocytes) in rAdVV vaccination (AZ and JJ), whilst increase in platelets and no change in monocytes in mRNA vaccination (PB, MD and CV). Overall decreased NK cells response was observed after rAdVV and mRNA vaccination. (3) Decreased CD4^+^ T cells activation, leukocyte chemotaxis and IL-18 signalling in rAdVV-based vaccination. Whilst in mRNA-based vaccination downregulation of different types of signalling pathways which are involved in proinflammation, profibrotic mediators and NFkB signalling pathways and positive regulation of NKT cell activation, platelet activation and aggregation, regulation of cell adhesion, cellular defence response, chemokine signalling pathways. (4) After 3^rd^ dose of COVID-19 vaccination (BP and MD) a strong humoral immune response was observed slightly higher than after the 2^nd^ dose of vaccination (mRNA COVID-19 vaccination only). Cellular response was robust, and decreased CD4^+^ TCM, NKT cells and a trend of increased Tregs were observed. (5) Cytokine CXCL-10, MCP1, IL-1RA were upregulated even after 3-6 months of COVID-19 vaccination (irrespective of the vaccination types). These cytokines could potentially be use as a biomarker of vaccine efficacy.

Our scRNA-seq data highlight that monocytes, CD4^+^ T cells, NKT, and NK cells predominantly changed depending on the type of COVID-19 vaccination. A recent study compared humoral and cellular immune response for 4 different vaccines and identified that Ad26.CoV2.S [Johnson & Johnson (JJ)] does not induce a strong immune response^57^. Similarly, our data also highlighted that JJ was not able to induce enough antibodies and antigen-specific T cells compared to mRNA and rAdVV (AZ) vaccines. Our results confirm other recently published studies especially for CD3^+^ T cells which were reduced after 2^nd^ and 3^rd^ dose of vaccination^58^, however, published data suggested persistent increase in the number of NK cells which is contrast with our study described in this report. We found that after the 2^nd^ vaccination (PoV2; mRNA PB), there was an increase in percentage of CD16^+^CD56^+^ NK cells, whilst after 6 months there was decrease in CD16^+^CD56^+^ NK cells which did not increase after the 3^rd^ dose of vaccination but again increased after the 4^th^ dose. A dynamic change in NK cells after mRNA vaccination (PB), whilst after rAdVV vaccination (AZ or JJ) there was decrease in NK cells after the 2^nd^ dose. This potentially alludes that NK cells migrate to the injection site or draining lymph nodes in vaccination individuals after COVID-19 vaccination^56^ and also depends upon sample collection after the vaccination. Furthermore, in case of cellular immunity, there is a strong and direct link among the vaccine-elicited T cell response and the capability of virus elimination^59–61^. Previous studies have provided enough support that those effective prophylactic vaccines against replicating viruses do engage strong cellular T cell-based immunity^62,63^. It is already known that adenovirus (Ad5-nCoV) does stimulate rapid T cell response^62,64^, however, the critical factors which are responsible for T cell-mediated immune protection against SARS-CoV-2 COVID-19 vaccines is still in his infancy. Our findings suggest that mRNA vaccination has strong spike-specific T cell response for mRNA COVID-19 vaccines (PB and MD), whilst rAdVV based vaccine (AZ and JJ) has reduced antigen-specific response after the 2^nd^ dose of vaccination. mRNA COVID-19 vaccines from CV were unable to induce antigen-specific response in 3-6 months post-vaccination and have reduced levels of IgG Spike S1 and RBD antibodies levels, respectively.

Earlier, it was reported that adenoviruses and SARS-CoV-2 mRNA vaccination activate several innate immune signalling pathways that result in the secretion of a number of pro-inflammatory cytokines^24,25,56,62^. Therefore, these pro-inflammatory cytokines could help us to predict effective immune cell stimulation and inform about induction of robust adaptive cellular immune responses. The results of these studies are consistent with our findings that immunization is associated with increased expression of pro- and anti-inflammatory cytokines, such as *MCP1 (CCL2), IL-1RA* and *CXCL-10* even after 3-6 month of vaccination. This result shows that T cells in vaccine recipients, unlike patients with critical COVID-19, are controlled and well ordered. Furthermore, cytokine CXCL-10, spike S1 and ageing correlations revealed that the ageing process affect the CXCL-10 production which could results in decline of protective antibodies. Thus CXCL-10 levels could be used as an important biomarker of vaccine efficacy. MCP1 (CCL2) was negatively correlated with RBD antibody levels in post vaccinated individuals (mRNA and rAdVV vaccines), implicating effective involvement of B cell immune response by innate immune system. Collectively, these findings help to illustrate the possible molecular basis of post-vaccination response, leading to a better understanding of the mechanisms of the T and B cell immune responses.

CD4^+^ TCM are specifically required for long-lived immunity in infectious disease such as influenza^65^ and these cell types increase after influenza vaccination. The major discovery based on cell type composition based on flow cytometry data showed a decrease in CD4^+^ TCM in mRNA after 2^nd^ and 3^rd^ doses of mRNA vaccination (PB) based on validation cohort. These results are in line with other recent study in which they described the reduced number of CD4^+^ TCM after 2^nd^ dose of mRNA vaccination^66^.

The scRNA-seq analysis of DEGs suggested for CD4^+^ NKT cells after rAdVV vaccination (AZ; PoV2) that several genes were upregulated (*GADD45B, IER2*) and that these gene are involved in modulation of Mcyobacterium *tuberculosis* (M. tb) host immune response, translation elongation, epigenetic changes and TCR downstream pathways as inferred from gene enrichment pathways (Suppl. Fig. 5 & 6). Recent animal model study also suggested that prior M. tb infection could generate protective immunity against severe COVID-19 disease^67^. Therefore, it implies that certain pathways must be common in NKT cells to generate immunity against bacterial and viral infections. The most common genes which were significantly downregulated (*FCGR3A, PTPRC, CD38, CD69, KLRB1*, and *IL2RB*) after rAdVV vaccination (AZ; PoV2) suggest that activation of leukocytes, natural killer cell mediated cytotoxicity, and neutrophil degranulation was hindered after vaccination in CD4^+^ NKT cells based on gene enrichment analysis. Analysis of DEGs in megakaryocytes before and after AZ (rAdVV) vaccination, *CD2* and *IGTB2* were upregulated. Pathway enrichment analysis suggested that purine ribonucleotide biosynthesis, regulation of NFkB signalling, and apoptotic signalling are involved. The downregulated pathways correlated with CD4^+^ T cell regulation and humoral immune response. Thus, there could be a nexus of platelets and T cells after post-vaccination (AZ) in some of the severe side-effects (vaccination induced thrombocytopenia), and they were related with dysregulated platelet activation. In our study, we found that platelets are inflammatory in nature after rAdVV vaccination whilst mRNA vaccination had no effect on platelet inflammation. Both types of monocyte populations (cMonocytes and pMonocytes) were decreased after AZ vaccination in numbers. Whilst, when pathway analysis was performed on these significantly changed genes and compared with PrV1, we found that the inflammatory response, response of bacterium, defence response to fungus and leukocyte cell-cell adhesion pathways were upregulated and that pathways related with cytokines (IL-2, IL-17, IL-18) were decreased.

Growing evidence suggests that NK cells may restrain potentially pathogenic effector T cell responses and that this immunoregulation may itself be regulated by CD244^52^. CD244 is expressed by all NK cells, γδ T cells, basophils and monocytes, and CD8^+^ T cells^48^. The ligand for CD244 is CD48 which increases under inflammatory conditions in particularly after exposure to type I & III IFN cytokines such as IFNα, IFNβ, and IFNγ on human peripheral blood mononuclear cells^50,52,68^. Notably, CD48 is also considered as a target of immune evasion by viruses, and this could be applicable in case of SARS-CoV-2 virus as earlier studies highlighted that during murine cytomegalovirus (CMV) infection, the mucin-like protein m154 reduces CD48 expression on macrophages, which limits NK cell-mediated control of viral infection^52^. Considering this evidence, we found that AZ (rAdVV) vaccine had reduced expression of CD244 on NK and NKT cells, whilst PB (mRNA) vaccine is still able to maintain the CD244 levels at mRNA and protein level. However, CD48 expression was somewhat different as AZ (rAdVV) vaccination had increased expression on both CD4^+^ TEM and TCM as well as CD8^+^ TCM. Whilst on PB mRNA vaccine had increased CD244 expression on CD4^+^ TCM and cMonocytes reflecting a potential distinct T cell response with different vaccines.

CD2 expression was very similar to CD48. CD48 binds to LFA3 (CD58) which is involved in the formation and organization of the immunological synapse between T cells and antigen-presenting T cells^51^. CD58 on the cell surface participates in potentiating effector-target adhesion during antigen-specific recognition^68^. Furthermore, CD2-CD58 interaction is also important for controlling the IL-12/IFN-γ positive feedback loop between monocytes and activated T cells^69^. CD2-CD58 interaction can induce non-proliferative Tregs with the production of large quantities of IL-10^68,70^. CD2 expression was high on cMonocytes in both AZ (rAdVV) and PB (mRNA) vaccination at protein levels compared with PrV1 state. Additionally, CD2 protein expression on CD8^+^ TEM I cells were low in AZ (rAdVV) whilst PB (mRNA) vaccination it was high. Thus, it appears that in AZ (rAdVV) vaccination antigen-specific response was less because of less effective communication among these co-stimulatory compared to mRNA vaccine (PB) which has effectively activated CD58-CD2 immune synapse formation for memory T cells and their effector functions. This agrees with our pathway analysis which revealed that after AZ (rAdVV) vaccination IL-10 signalling pathway was less active.

The C-C chemokine receptor 5 (CCR5)/Rantes pathway was altered in critical COVID-19 patients, and reduction in CCR5 was suggestive for improved out come in the patient recovery as was less viral load^71^. Furthermore, large scale genome wide studies suggested that CCR5 is linked with COVID-19 severity^72–74^. Reduced CCR5 expression and immunosuppression was found in long-term COVID-19 syndrome (LTCS) patients and increase in expression of *CCR5* by treatment with leronlimab was correlated with an increase in T cells and a decrease in IL-10 and C-C chemokine ligand-2 (CCL-2)^75^. In our study, we found less abundance of CCR5 in rAdVV vaccination (AZ) whilst in PB (mRNA) vaccination it was not changed compared with PrV1 individuals. However, how this does help the COVID-19 vaccination response and outcome is still unclear and further studies are required to understand the role of CCR5 in COVID-19 disease. More interestingly, another key finding was differential expression of MALAT1 for AZ (rAdVV) and PB (mRNA) vaccination. We found that the lncRNA MALAT1 was upregulated like CD48 expression in AZ compared with PrV1 or PB (mRNA) vaccination. Thus, we believe that it could be positively regulating the expression of CD48. MALAT1 is involved in memory T cell generation. A previous study by Kanbar and coworkers suggested that Malat1 groups with *trans* lncRNAs that exhibit increased RNA interactions at gene promoters and gene bodies^76^. Moreover, Malat1 was also associated with increased H3K27me3 deposition at a number of memory cell-associated genes through a direct interaction with Ezh2, thereby promoting terminal effector and t-TEM cell differentiation^76^. MALAT1 is downregulated in tissues from patients with multiple sclerosis and mice with experimental autoimmune encephalomyelitis and knockdown of MALAT1 promoted Th1/Th17 polarization and inhibited T regulatory cell differentiation *in vitro*^77^. Furthermore, MALAT1 suppression is a hallmark of CD4^+^ T cell activation and controls IL-10 expression in Th cells^78^. Keeping this line of evidence, we suggest that higher expression of MALAT1 after AZ vaccination (rAdVV) could be affecting the proper CD4^+^ and CD8^+^ T cells activation compared with mRNA vaccination (PB), therefore MALAT1 could be affecting the antigen-specific immune response. Our data suggested that AZ (rAdVV) vaccination have less antigen-specific IFN-γ producing cells compared with PB mRNA vaccine. However, further studies are needed to understand how different COVID-19 vaccinations influencing different lncRNA and epigenetic changes could affect the antigen-specific T cells which could help to explain the long-term protection of vaccination as antibodies levels decline after 6 months of vaccination. Therefore, importance of antigen-specific T response is the key for the future research after vaccination.

In sum our data highlighted that different COVID-19 vaccine platforms have distinct mechanism of actions to generate vaccine-specific immunity. In future, data generated using scRNA-seq will help us to access durability of the vaccine protection and guide future vaccination strategies to delineate the optimal outcomes against emerging SARS-CoV-2 variants.

### Limitation of the study

Our study has several limitations. Due to limited numbers of AZ, CV and JJ vaccinated individuals in our cohort, it is insufficient to confirm the broader effects of vaccination on immune response. Furthermore, volunteers were free to choose second or third choice in real-world data setting, therefore, for the 3^rd^ dose of vaccination, we had mostly PB or MD (mRNA) COVID-19 vaccination and no rAdVV vaccine samples have been available. Thus, a direct comparison for rAdVV and mRNA vaccination was not possible for third dose of the samples. Nevertheless, extension of these studies also with novel upcoming vaccines against specific SARS-CoV-2 variants will help to decide on the frequency and combination of vaccines for most effective SARS-CoV-2 disease protection.

## Acknowledgements

We are grateful to all the volunteers for donating blood samples for the study. We thank Dr Denis Witt, Dr Lisa Ruisinger, Miss Anna Liu, Mr Surya Pal Sekhar, Mr Omer Khalid for technical assistance with blood sample collections and PBMCs isolation. Prof Dr Sara Y Brucker for infrastructure support for the experiments. We also acknowledge UKT, FACS core facility for providing excellent infrastructure for flow cytometry measurements and technical assistance. We gratefully acknowledge Prof. Huu Phuc Nguyen, Ruhr-Universität Bochum, Germany for critical reading of the manuscript.

## Funding

This project was supported by funded by the Deutsche Forschungsgemeinschaft (DFG Project no 286/2020B01 – 428994620) to OR. NGS sequencing methods were performed with the support of the DFG-funded NGS Competence Center Tübingen (NCCT) (INST 37/1049-1). This project was analysed using the de.NBI cloud infrastructure (DFG Project no 031A537B, 031A533A, 031A538A, 031A533B, 031A535A, 031A537C, 031A534A, 031A532B). CR was supported by fortüne/PATE (no. 2536-0-0/1) from the medical faculty, University of Tübingen. YS was supported by fortüne funds (no. 2646-0-0) from the medical faculty, University of Tübingen.

## Conflicts of Interest

The authors declare no conflict of interest. The funders had no role in the design of the study; in the collection, analyses, or interpretation of data; in the writing of the manuscript; or in the decision to publish the results.

## Author’s contribution

YS: Conception of the project and overall project management, performed most of the experiments (Single cell RNA-seq, flow cytometry) and data analysis (single cell-RNA-seq and all the flow cytometry data) and writing the manuscript

ASS: Performed single cell RNA library prep and sequencing

GG, VH: Single cell data analysis and figure preparation and writing materials and methods.

UR, RB, MA: Performed the PBMCs isolation and antibodies assays, multiplex cytokine assays, flow cytometry staining

EBA: Single cell RNA sequencing and data preparation

CR, MK: Sample collection, PBMCs isolation and data analysis JHS: Cohort organization and metadata collection

PK: Sample collection, data discussion

MSS, NC, SO: Project conception, experimental planning, data discussion and funding acquisition

DA, SN: Data discussion and editing the manuscript

OR: Project conception, experimental planning, data discussion and funding acquisition and writing the manuscript

DeCOI: Provided tools for the single cell seq analysis and data discussion

## Data availability

scRNA-seq GEX data is available on EGA accession no. XXXXX. Single-cell gene expression and feature barcoded antibody processed data will be available on the FastGenomics database (Seurat_objects_Singh_COVID19_VAC). Raw single cell data (fastq files) are available from the corresponding author upon mutual reasonable request.

## Notes

### Competing Interest Statement

The authors have declared no competing interest.

