## Supplementary material for "Comparative multi-OMICS single cell atlas of five COVID-19 (rAdVV and mRNA) vaccines describe unique and distinct mechanisms of action": Suppl.info

[1]Institute of Medical Genetics and Applied Genomics, Eberhard Karls Tübingen University, Tübingen, Germany.
[2]NGS Competence Centre Tübingen (NCCT), Eberhard Karls Tübingen University, Tübingen, Germany.
[3]Department of Women's Health, Research Institute for Women's Health, Eberhard Karls Tübingen University, Tübingen, Germany.
[4]Quantitative Biology Center (QBiC), Eberhard Karls Tübingen University, Tübingen, Germany.
[5]Centre for Neurology, Hertie Institute for Clinical Brain Research, Eberhard Karls Tübingen University, Tübingen, Germany.
[6]Institute of Tropical Medicine, Eberhard Karls Tübingen University, Tübingen, Germany.
[7]Faculty of Medicine, Department of Immunology and Inflammation, Imperial College London, London, UK
[8]DeCOI members are presented in https://decoi.eu/members-of-decoi/

*Lead Contact

Dr Yogesh Singh
Institute of Medical Genetics and Applied Genomics
NGS Competence Centre Tübingen (NCCT)
University of Tübingen
Calwerstraße 7, 72076, Tübingen
Germany


Key words: COVID-19 vaccines, PBMCs, NK cells and cytotoxic cells

Running title: Single cell sequencing of COVID-19 vaccinated individuals

**Suppl. Information**

### Materials and Methods

#### *Human subjects, experimentation, and Ethics statements*

110 healthy volunteers were recruited for the study under informed consent and a baseline health questionnaire was also completed for their vaccination intake. The study was approved by the Institutional Review Board of the University Hospital Tübingen and Eberhard-Karls University of Tübingen (Ethik-Kommission and der Medizinischen Fakultät der Eberhard-Karls-Universität and am Universitätsklinikum Tübingen, Germany, Ethics no: 355/2021BO2). The study was conducted within full compliance of Good Clinical Practice as per the Code of German Regulation. The study was registered at clinicaltrials.gov (NCT04873128). All studies adhere to the principles of the Declaration of Helsinki, and written informed consent was obtained from every participant, patient, or legal representative.

We used two cohorts for the study, first the discovery cohort (baseline and after vaccination; n=28; n=3-5 per vaccine except SP, where n=1) and the second the validation cohort (vaccinated follow up 6-12 months; n=82) was used due to limited number of pre-vaccinated individuals available to participate in the study (Supp. Table 1: Patient demographics). In discovery cohort [pre-vaccinated (PrV1), post-vaccination 1 (PoV1), pre-vaccination 2 (PrV2) and post-vaccination 2 (PoV2)], we performed Spike S1 IgG antibody detection, Spike-specific T cell detection, sc-RNA-seq and 14-colour immunophenotyping. In validation cohort [post-vaccinated 3- month (PoV3), pre-vaccination 3 (PrV3), post-vaccination 3 (PoV3) and post-vaccination 4 (PoV4)], we analysed 3-6M PoV2 and PoV3 Spike S1 IgG antibody, 13-different cytokines which are involved in cytokine storm and 14-colour immunophenotyping.

#### *PBMC and Plasma isolation*

S-Monovette 92x16 mm 9ml AH $NH4^+$Heparin: 16 I.U./ml tubes (#02.1064; Sarstedt, Germany) were used to collect 8-9ml sample from pre-vaccinated and post-vaccinated individuals. Blood samples were left in an upright position for 1-2 hours before isolation of PBMCs at room temperature so that all the samples can be processed at the same time. The plasma layer was carefully removed, transferred to a conical vial, spun at 2000 RCF for 10 minutes, and the supernatant transferred to microtubes in 1 mL aliquots. Plasma was stored at -80°C until further use.

The remaining blood was diluted 1:1 with DPBS (#D8662, Sigma, Germany) and carefully added to 50mL of Pancoll human (density1.077g/ml) (#P04-60500, PAN Biotech, Germany) and spun at 400 RCF for 20 minutes at room temperature without breaks. After centrifugation, PBMCs were carefully taken and transferred to a new 50mL of Falcon tube and added 20mL of PBS to remove all the remaining Pancoll by centrifuging the tubes at 400 RCF for 5 minutes at room temperature. PBS was discarded, and the pellet was re-suspended in again 2-3mL of PBMS and cells were counted with TC20 Automated Cell Counter (BioRad, Germany). Cells were again pelleted by centrifugation at 400 RCF for 5 minutes and resuspended in heat-inactivated FBS (Thermofisher, Germany) containing 30% DMSO (Sigma-Aldrich) at a concentration of 3-5 million cells/mL in 2.0 ml cryovials (#121263, Grenier Bio-One, Germany). Cryovials were frozen overnight to -80°C using freezing containers (Corning® CoolCell™ LX Cell Freezing Container) and cells were kept until use in -80°C for short-term storage.

**Single cell RNA sequencing (sc-RNA-seq)**

**Antibody staining:** Cryovials were taken from the storage (-80 $^{0}$C freezer and kept on dry ice if many samples were processed at the same time) and immediately thawed in water bath at 37 $^{0}$C for 2-3 minutes (until a small ice crystal remain). In a biosafety hood, slowly transferred thawed cells to a 50 ml of conical tube using a wide-bore pipette tip. The cryovial was rinsed with 1 ml of warm complete RPMI1640 medium (RPMI1640 with glutamax+10%FBS+1%Anti/Anti) (all from Thermofisher, Germany) and added the rinse dropwise (1drop per 5 seconds; to remove DMSO from frozen PBMCs) to 50 ml of conical tube whilst gently shaking the tube. All the cell washes were performed at room temperature (RT). Further, sequentially, cells were diluted in 50 ml of conical tube by incremental 1:1 volume additions of media for a total of 5 times with 1 minute wait between the additions. Complete RPMI1640 medium was added a speed of 1 ml/3-5 second to the tube whist swirling the tube by another hand (total medium 30-32 ml). After adding the complete medium, cells were centrifuged at 400 RCF for 5 minutes at RT, after centrifugation, most of the supernatant was removed leaving approximately 1 ml of medium. Cell pallet was resuspended in this volume using wide bore pipette tips and added additional 9 ml of complete RPMI1640 dropwise whilst swirling the tube with another hand and cells were centrifuged again at 400 RCF for 5 minutes. Before adding the complete medium 20 µl cells were taken for the tube for counting the cells using Trypan blue Automated Cell Counter method (BioRad). A minimum cell viability of 80% was considered for single cell sequencing. After centrifugation cells were resuspended into $0.25x10^{6}$ ml cell staining buffer (#420201; BioLegend, San Diego, USA) or 1xPBS+2.0% BSA and transferred to an Eppendorf tube. Cells were washed 3x with cell staining buffer to remove any residual serum from the tube. Before staining the cells with TotalSeq™ Universal Cocktail (TotalSeq™-C, #399905, BioLegend, San Diego, USA), the lyophilized panel vial was equilibrated at RT for 5 minutes and spun down at 10,000 RCF for 30 seconds at RT. Lyophilized antibody cocktail was rehydrated by adding 52 µL of cell staining buffer, vortexed for 30 seconds and left for 5 min incubation at RT. The reconstituted vial was vortexed again and spun down at 14,000 RCF for 10 min at 4 $^{0}$C. Whilst centrifugation of antibody cocktail was ongoing, cells were blocked by adding 2.5 µL of Human TruStain FcX™Fc blocking reagent (#422302; BioLegend, San Diego, USA) in 22.5 µL cell staining buffer. Cells were left for incubation at 4 $^{0}$C (keep the cells on ice) for 10 min in an Eppendorf tube. After 10 minutes of incubation, 25 µL of reconstituted cocktail was added to the cells together with 25 µL of cell staining buffer and Human TruStain FcX™Fc blocking reagent (total volume 50 µL) and left for 30 minutes incubation at 4 $^{0}$C (keep the cells on ice). After antibody incubation, cells were washed at 400 RCF at RT for 4x to remove any residual volume of unbound antibodies and suspended in 200-250 µl of cell staining buffer to achieve optimal 700-1200 cells/µl for 10x Chromium Single experiments (all the samples kept on ice after this step until cells were loaded on 10x Chromium Chip).

**10x Chromium single cell cDNA preparation:** Before loading the cells on the 10x Chromium Chip, cells were counted using Acridine orange and PI dye for live and dead cells. In the most of the sample's cells were >90% viable whist in few samples it was 76% viable cells were used. We used Chromium Next GEM Single 5' Library and Gel Bead Kit V1.2, 16 rxn PN-1000165. Gel Beads-in-emulsion (GEM) were generated by combining barcoded single cell VDJ 5' Gel Beads, a master mix with cells (30,000) and portioning oil on Chromium Next GEM Chip G. Immediately following GEM generation, the Gel Beads dissolved, and any co-partitioned cell were lysed. Oligonucleotides containing (i) an Illumina R1 sequence (read 1 sequencing primer), (ii) a 16 nucleotide 10x Barcode, (iii) a 10-nucleotide unique molecular identifier (UMI), and (iv) 13 nucleotide template switch oligo (TSO) were released and mixed with cell lysate and master mix containing reverse transcription (RT) reagent and poly(dT) RT primers. In the cell lysate and the released Gel Bead primer incubated with the Master Mix containing RT reagents, produce10x barcoded, full length cDNA from poly-adenylated mRNA. Simultaneously, in the same partition, the Gel Bead captures the cell surface protein Feature Barcode containing (i) a Nextera Read 2 (Read 2N), (ii) a 15 nucleotides Feature Barcode, and (iii) Capture Sequence. Incubation of the GEMs with the master mix containing RT reagents, produce 10x Barcoded, DNA from the cell surface protein Feature Barcode.

GEMs were broken and pooled after GEM-RT reaction mixtures were recovered. Silane magnetic beads were used to purify the 10x Barcoded first-strand cDNA from the post GEM-RT reaction mixture (to remove the biochemical reagents and primers). After cleaning up, target enrichment was performed from first-strand cDNA for TCR and BCR library preparation. Further, 10x Barcoded, full length cDNA from poly-adenylated mRNA and DNA from protein Feature Barcode were amplified. Amplification generates sufficient material to construct multiple libraries from the same cells such as T or/and B cell Enriched libraries, 5' Gene Expression libraries and Cell Surface Protein libraries. Amplified cDNA from poly-adenylated mRNA and the amplified DNA from cell surface are separated by size selection for generating V(D)J and 5' Gene Expression and Cell Surface libraries.

**Target enrichment (TCR and Ig) from cDNA and library construction:** Amplified full-length cDNA was used for enriching full length V(D)J (10x Barcoded) via PCR amplification with primers specific to either TCR or Ig constant regions. both T and B cells were present in the PBMCs, therefore, TCR and Ig transcripts were enriched in separate reaction from the same amplified cDNA and P5 was added during enrichment PCR reaction. Enzymatic fragmentation and size selection were used to generate variable length fragments that collectively span the V(D)J segments of the enriched TCR and Ig transcripts prior to library construction. P7, a sample index, and an Illumine R2 sequence (Read 2 primer sequence) were added via End Repair, A-tailing, Adaptor Ligation, and sample Index PCR. The final libraries contained the P5 and P7 priming sites used in Illumina sequencing.

**5' Gene Expression (GEX) library construction:** Amplified full-length cDNA from poly-adenylated mRNA was used to generate 5' Gene Expression library. Enzymatic fragmentation and size selection were used to optimize the cDNA amplificon size prior to 5' gene expression library construction. P5, P7, a sample index, and Illumina R2 sequence (Read 2 primer sequence) were added via End Repair, A-tailing, Adaptor Ligation, and sample Index PCR. The final libraries contained the P5 and P7 priming sites used in Illumina sequencing.

**Cell Surface Protein (ADT or feature bar code antibodies) library construction:** Amplified Feature Barcode was used to construct Cell Surface Protein library. P5, P7, a sample index, and Nextera Read 2 (Read 2N primer sequence) were added Sample

Index PCR. The final libraries contained the P5 and P7 priming sites which were used for Illumina sequencing.

All the libraries were pooled and sequenced on an Illumina HiSeq 4000, in a S1 flow cell, using the recommended sequencing read lengths of 28 bp (read 1), 8 bp (i7IndexRead) and 91 bp (read 2). 30,000 sequence reads were obtained for the GEX libraries, 5000 reads for TCR and Ig libraries and 10,000 reads for Feature Barcode (FB) Protein per cell).

**Single cell data processing:**

**Processing of raw sequencing reads:** Cell Ranger (v.6.0.2, 10xGenomics) was used to demultiplex raw sequencing data (cellranger mkfastq) and create FASTQ files. To perform alignment, filtering, barcode and UMI counting of the GEX and FB libraries as well as processing of the TCR and Ig libraries Cell Ranger (v.6.0.2) was employed (cellranger multi pipeline). The Ensembl reference GRCh38.p13 release 98 (corresponding to pre-built indices provided by 10x genomics 2020-A) was employed for read mapping.

**Sample integration and clustering:** The Seurat package v4.0.6 was used for downstream sample integration and data analysis. Cells were removed out which had less than 200 genes or more than 10% mitochondrial counts to exclude dead or dying cells from the analysis. Additionally, cells with more than 2500 genes were excluded to avoid multiplets. After filtering, 329,920 cells remained.
Data was normalized on a per-sample basis with Seurat's "*LogNormalize"* method. The 2000 genes with the hightest variance to mean ratio for each sample were identified with the "*FindVariableFeatures"* function. To remove subject batch effects, samples were integrated (Seurat's "*IntegrateData"*) according to integration anchors (genes) conserved across the samples identified using the "*FindIntegrationAnchors*" function with reciprocal principal component analysis (PCA) with 50 dimensions.
Then, on the integrated Seurat object, a PCA was perfomed and the first 50 components were used to for the Shared Nearest-neighbor (SNN) graph construction with the "*FindNeighbors*" function. Modularity optimization was performed on the SNN graph with the "*FindClusters*" function, using the Louvain algorithm with a resolution of 0.4 leading to 27 clusters. The function "*FindAllMarkers"* was employed to identify gene markers for each cluster. The antibody-derived tags data was normalized across cells with a center log ration transformation using Seurat's "*NormalizeData*" with the "*CLR"* method. Scaling was performed with the "*ScaleData*" method.
Cell type annotation was performed manually by using the list of markers provided in XXX and relevant surface proteins (130 feature barcoded antibodies). The annotations were verified with Seurat's Azimuth PBMC reference. Cell clusters were removed if less than 50 cells.

Cell annotation:

We processed all the samples and clustered and annotated according to the scRNA-seq expression profiles using Seurat[1] (detail in material and methods). Gene per cell and mitochondrial content were filtered out (Suppl. Fig. 3, 4). Principal component analysis (PCA) guided UMAP analysis was performed. In total, 27 clusters were obtained based on 50 principal components and test resolution equivalent to 0.4 when used unsupervised clustering and cell clusters were plotted in UMAP space (Fig. 2b). Each cluster was manually assigned to a specific cell type based on the expressed gene listed in Suppl. Tables 2&3 (Fig 2b, c). Further, we used Azimuth package to predict the cell type for each of the cells that were found with unsupervised clustering (Fig. 2b,c). In majority of the cases, manual gating was matched for most of the cell cluster. Therefore, we followed the Azimuth-based cell annotation for this study to define the cell types for each cell cluster (Suppl. Fig. 6). The majority of the cells belong to CD4$^{+}$ T naïve (cluster 0; CD4$^{+}$ TN) and helper T cells (IL-7R$^{+}$ CD3E$^{+}$ cells; 5 clusters IL-7R$^{+}$ T4 cells 1, 10, 17, 22, 15), and cluster 15 specifically belongs to regulatory T cells (Tregs) (cluster 15 marker genes; FOXP3$^{+}$, IZKF2$^{+}$, IL-2RA$^{+}$, RGS1$^{+}$ and CTLA-4$^{+}$)[2]. IL-7R$^{+}$ cells were further divided in to CD4$^{+}$ TN, T effector memory (TEM) CD4$^{+}$ or T central memory (TCM) CD4$^{+}$ cells based on CCR7, SELL, AQP3, CD27, CD28 mRNA or feature bar code antibody (protein) expression (Suppl. Table 1; Cluster genes)[3]. However, most of TCM and TEM CD4$^{+}$ T cells contain a mixture of genes that are difficult to distinguish among two specific cell subsets based on common gene markers (Suppl. Fig. 5, Suppl. Table 4; cluster gene). Overall, based on unsupervised clustering, we can identify the 25 major cells clusters (confidently) based on differential gene expression.

CD8$^{+}$ T cells (clusters 4, 5, 6, 8), NK cells (NCAM1$^{+}$ or CD56; clusters 2, 18, 19) (Fig. 2c and Suppl. Table 4; cluster gene), CD8$^{+}$ NKT cells (cluster 11; CD3E$^{+}$ GZMH$^{+}$, KLRD1$^{+}$ PRF1$^{+}$, FCGR3A$^{+}$ and NCAM1$^{+}$), and B cells (clusters 7, 9; MS4A1$^{+}$ or CD20 or CD19; MS4A1$^{+}$ B cells ) belong to lymphoid cell compartment, whilst classical monocytes (cluster 3; cMonocytes - LYZ$^{+}$, CD14$^{+}$, S100A9$^{+}$, FCN1$^{+}$, VCAN$^{+}$, IL1B$^{+}$), plasmacytoid monocytes (cluster 12; pMonocytes - CD16a or FCGR3A), dendritic cells (clusters 20, 21; DCs - ITGAX or CD11c), and megakaryocytes/platelets (cluster 16; PPBP$^{+}$ cells) made separate clusters of myeloid cell compartment (Fig. 2b, c). Similarly, like CD4$^{+}$ T cells, CD8$^{+}$ T cells were further classified into TN, TEM, and TCM based on PRF1, GZMK, GZMH, GNLY, CD8A, CD69, CD63, CCR7, and PTPRC marker expression (Fig 2b, c). Furthermore, CD8$^{+}$ T cell clusters also contain gamma-delta (γδ)-T cells (cluster 14) and mucosal invariant T cells (MAIT; cluster 13) in the CD8$^{+}$ T cell compartment. NK cells were characterized by PRF1, NKG7, FCGR3A, CX3CR1, KLRD1, IL2RB, and PTGDS markers (Suppl. Table 4; cluster gene). Furthermore, B cells clusters were identified with IGHM, MS4A1, BANK1, CD79B, JCHAIN, BANK1, and BLK markers for different subsets of B cells (Suppl. Fig. 5b), whilst DCs (FCER1A$^{+}$DCs I and II) were identified using FCER1A, CD1C, HLA-DRA, HLA-DQA1, HLA-DQB1, PTGDS, CCDC50, PLD4, IRF8, and TLR9 markers (Suppl. Table 4; cluster gene). The classical monocytes were classified based on S100A8/9, FCN1, VCAN, LYZ, IL1B, TREM1, NLRP3, DUSP6, and HLA-DQB1/DQA1, and plasmacytoid monocytes had CXCL16, MS4A7, FCGR3A, OAZ1, PRDX1, as well as C1QA higher marker gene expression (Suppl. Table 4; cluster gene). Megakaryocytes/platelets were defined based on PPBP, NRGN, PF4, TUBB1, CAVIN2, SNCA, GP9, C2orf88, ITGB3, TREML1, SPARC, and F13A1 marker genes (Suppl. Table 4; cluster gene). Overall, based on unsupervised clustering, we can identify the 25 major cells clusters based on differential gene expression.

**Cell type proportions analysis:** the number of cells within a cell type was normalized by the total number of cells in each sample. The cell proportions were plotted in a boxplot by vaccination status. Significant differences among groups were determined by the Wilcoxon rank-sum test.

**Differential gene expression analysis:** Seurat's function "FindMakers" was employed to determine differentially expressed genes for each of the cell types among the different vaccination groups. The test used to determine differential gene expression was the default Wilcoxon rank test, with Bonferroni multiple-test correction. A log2FC threshold of 0.5 was applied, and a significance threshold was set to an adjusted p-value of <=0.05.

**Metascape analysis*:***

*Pathway and Process Enrichment Analysis*
For each given gene list from the specific cell cluster (differentially regulated genes either upregulated or downregulated gene were used separately for pathway analysis), pathway and process enrichment analysis has been carried out with the following ontology sources: KEGG Pathway, GO Biological Processes, Reactome Gene Sets, Canonical Pathways, Cell Type Signatures, CORUM, TRRUST, DisGeNET, PaGenBase, Transcription Factor Targets, WikiPathways, PANTHER Pathway and COVID using online metascape tool (https://metascape.org)[4]. All genes in the genome have been used as the enrichment background. Terms with a p-value < 0.01, a minimum count of 3, and an enrichment factor > 1.5 (the enrichment factor is the ratio between the observed counts and the counts expected by chance) are collected and grouped into clusters based on their membership similarities. More specifically, p-values are calculated based on the accumulative hypergeometric distribution, and q-values are calculated using the Benjamini-Hochberg procedure to account for multiple testing. Kappa scores are used as the similarity metric when performing hierarchical clustering on the enriched terms, and sub-trees with a similarity of > 0.3 are considered a cluster. The statistically most significant term within a cluster was chosen to represent the cluster[4].

**Flow cytometry staining and data analysis:**

*14-colour immunophenotyping*
A total of 1–2 x$10^6$ PBMCs per volunteer per time point were used for 14-colour flow cytometry panels (Suppl. Table 2: antibody Panels). PBMCs were first resuspended with Human TruStain Fcx (Biolegend) for 10 minutes at room temperature and then stained with the following antibodies as mentioned in Table X. In brief, firstly, to distinguish between live from dead, the cells were incubated with LIVE /DEAD Fixable Infra-Red Dead stain (Thermofisher) for 15 min at room temperature (RT) into 1:40 diluted dye in DPBS. Subsequently, after LIVE/DEAD staining, cells were stained with surface markers in DPBS (Sigma) with Super Bright stain Buffer (Thermofisher) for 30 min at RT. After antibody staining, cells were washed once (600 RCF for 5 minutes at RT) and fixed (Fix/Perm buffer 1x) for 30-45 minutes and washed 2x with DPBS and resuspended in 100ul DPBS and samples were acquired on flow cytometry either same day or next day. For each sample, 200,000 cells were acquired using BD LSRFortessa (core facility) equipped with 4 lasers (violet, blue and yellow-green and Red). Data were analysed using Flow Jo (Tree Star) and fluorescence minus one control (FMO) were used for setting up the arbitrary gates for the major cell markers initially as described earlier[5].

*Activation induced marker (AIM) detection, antigen-specific response after Spike-S peptide by means of cytokine detection using flow cytometry*

A total of 1–2 x$10^6$ PBMCs per volunteer per time point were used for culturing and added CD40 antibody (1ug/1x$10^6$ cells) for overnight before staining for LIVE/DEAD and AIM markers. In another experiments, 1–2 x$10^6$ PBMCs per volunteer per time point were used for culturing and added 1ug PepTivator®SARS-CoV-2 Prot_S, research grade/1x$10^6$ cells (#130-126-700; Miltenyi Biotec) for overnight. Next day cells were treated with ionomycin (0.5ug/ml) and Brefeldin A (1ug/ml) for 4 hours before staining for LIVE/DEAD and cytokine staining (Suppl. Table 2: antibody Panels). Surface staining was performed as described in preceding section until the fixation step. Before IC staining, cells were fixed for 30–45 min and permeabilized for 5 min followed by IC antibody incubation for additional 30 min at RT. Cells were washed (600 RCF for 5 minutes at RT) and resuspended in DPBS containing 2%FBS. Fixing of cells was performed irrespective of whether the panel was used for IC staining or not to prevent the possible contamination during the acquisition of the samples. All the staining procedures were performed in the dark to avoid the photo-bleaching of dyes. For each sample, 200,00-500,000 cells were acquired using BD LSRFortessa. Data were analysed using Flow Jo (Tree Star) and fluorescence minus one control (FMO) were used for setting up the arbitrary gates for the major cell markers initially as described earlier[5].

Data were analysed for normal percentage of the cells as well as using dimension reduction method (UMAP) in Flowjo software using default setting except for minimum distance (0.2 instead of 0.5) and population (n = 15) as advised by a plug-in. First, dead/debris was removed by gating (FSC-A vs SSC-A; linear scale) then using FSC-A vs FSC-W, we focussed on singlets. The singlets were again gated for live and dead discrimination. An equal number of live cells from each sample (PrV, PoV1, PoV2, PrV3, PoV3 and PoV4) were concatenated and exported as a single FCS file. This FSC file was subjected to UMAP analysis, each cell population was either monocytes or lymphocytes were again subject to subsequent UMAP analysis for clustering of specific sub-populations. Based on gating or antibodies-stained cells (data-driven) analysis was performed as mentioned in Fig.1 & 5.

***Anti-Spike protein S1, anti-spike protein RBD and anti-Nucleocapsid IgG antibodies multiplex flow cytometry ELISA***

To measure the amount of IgG antibodies against Spike protein S1, Nucleocapsid and Spike protein RBD in the plasma of vaccinated volunteers, we used SARS-CoV-2 Serological IgG panel (3-Plex), LEGENDplex™ Multi-Analyte Flow Assay kit (#741132; BioLegend, San Diego, USA). It is a bead-based multiplex assay utilizing fluorescence-encoded beads suitable for use on flow cytometry and followed the manufacture's instructions. In brief, we diluted plasma samples to 800-fold using serial dilution method, first diluted the samples 100-fold (e.g. 1ul of plasma sample with 99 ul of LEGENDplex™ Assay Buffer and secondly serially diluted 20 ul of that mixture into 140 ul of LEGENDplex™ Assay Buffer. After dilution of plasma, 8 standards were prepared using serial dilution methods (1:4 dilution). Prior to use, lyophilized SARS-CoV-2 Serological Panel Standard were reconstituted with 250 ul LEGENDplex™ Assay Buffer. Standard were mixed allowed the vial to sit at room temperature for 15 minutes and transferred to a C7 labelled Eppendorf tube and used as a top standard.

Standard was serially diluted to six more microcentrifuge tubes labelled as C6, C5, C4, C3, C2, C1 respectively. In each tube 75ul of Assay buffer was added. To prepare 1:4 dilution of the top standard by transferring 25 ul of the top standard C7 to the C6 tube and mixed well. In the same manner, C5-C1 samples were serially diluted. C0 was made with only Assay Buffer. All the reagents allowed to sit at room temperature for 30 minutes. For measuring the amount of antibodies levels, V-bottom 96-well plate was used. First, 16 wells were used for the standards (C7-C1 and C0; blank) and rest 80 wells used for the samples. In all the well, 25 ul of Assay Buffer was added. In the standard wells (C7-C0), 25 ul of each standard was added in duplicate and in the sample well 25 ul of 800-fold diluted plasma was added (single reaction). After adding the standard and samples, beads were vortexed for 30 – 60 seconds and 25 ul of beads were added to each well (total volume 75 ul) and beads were intermittently mixed while adding to the wells. Plate was sealed with a plate sealer and entire plate was covered with aluminium foil to protect the beads exposed to light. Plate was incubated for 2 hours at room temperature at 800 rpm on a plate shaker. After incubation, plate was centrifuged at 250 RCF for 5 minutes, using a swinging bucket rotor with microplate adaptor. Immediately after centrifugation, supernatant was dumped into a clinical waste biohazard container. The 96-well plate was blotted on a stack of clean paper towel and drained the remaining liquid from the well as much as possible without disturbing the beads. Washing step was performed by dispensing 200 ul of 1x wash buffer into each well and incubated the plate at room temperature for at least 1 minute followed by centrifugation at 250 RCF for 5 minutes. After centrifugation, wash buffer was discarded, and plate was dabbed on the pile of tissue paper to remove any excess wash buffer and then 25 ul of detection antibodies was added to each well. Plate was sealed with new plate sealer and covered with aluminium foil and kept the plate on a plate shaker at 800 rpm for 1 hour at room temperature. After incubation period, 25 ul of SA-PE antibody was added to each well. Again, the plate was sealed with a new plate sealer and kept on a plate shaker at 800 rpm for 30 minutes at room temperature. After incubation, the plate was washed 2x as described above and added 150 ul of 1x Wash Buffer to each well. All the experimental and washing steps were performed at RT. Beads were resuspended by pipetting or vertexing just before acquiring the samples on flow cytometry. Data analysis was performed using Biolegend's LEGENDplex™ data analysis software. Dilution correction was added during analysis. The concertation of each IgG antibodies was presented in ng or ug/ml and calculated with standard curve method.

**Detection of cytokines using 13-plex flow cytomix cytokines assay**
The levels of blood plasma cytokines and chemokine patterns were measured by a Multi-Analyte Flow Assay Kit, COVID19 Cytokine Storm panel 1 (13-plex) w/VbP (BioLegend, San Diego, USA). The magnetic beads-based multiplex assay was used in state of a traditional ELISA because of the higher sensitivity and broader dynamic range for quantitative measurement of 13 human inflammatory cytokines and chemokines including IL-6, CCL2 (MCP-1), G-CSF, INF-a2, IL-2, IFN-γ, IL-7, IL-1RA, CXCL10 (IP-10), CCL3 (MIP-1a), IL-10, TNF-a, CXCL8 (IL-8). In principle, the beads will be differentiated from each other based on their size and internal fluorescence intensities on a flow cytometer platform. Each bead set is bound with a specific antibody on its surface and forms capture beads for that analyte. Blood samples were drawn from healthy individuals who received mRNA-based vaccines at different time points at the institute of the medical genetic, university of Tübingen. We next analyzed their plasma for cytokine signatures by the Multi-Analyte Flow Assay platform. In the

standard preparation section: 1:4 dilution of the top standard (C7) was first prepared as the highest concentration, then serial dilution was done for C6, C5, C4, C3, C2, and C1 by taking 25 µl of diluted standard and added into 75 µl assay buffer. Following, 15µl of serum samples were equally diluted with 15µl assay buffer. Next, 25µl of the diluted samples was carefully transferred to each well. 25µl of mixed beads was added to each well. Importantly, beads were mixed well before using by vortex for 30 seconds to avoid bead settling. The plate was sealed with a plate sealer, covered entirely with aluminium foil to protect the plate from light, and put on a plate shaker at 800 rpm for 2 hours incubation at room temperature. After incubation, the plate was centrifugated at 250 RCF for 5 minutes, immediately the supernatant was carefully discarded by flicking the plate in one continuous and forceful motion and the plate was washed by 200µl 1X wash buffer. Following 25µl of detection, antibodies were added to each well, the plate was again sealed with a plate sealer, covered entirely with aluminium foil, and incubated for 1 hour at RT. After incubation, 25µl of SA-PE was directly added to each well without washing the plate and the plate was sealed and covered in the same manner as described in a previous step. Following the plate was centrifuged for 5 minutes and washed in the same manner as described before. Finally, 150µl of 1X wash buffer was added to each well and the samples were stored in the cold room until the reading day by flow cytometer.

**Statistics**

Cytometry data were analyzed using FlowJo 10.8.1. Statistical analyses were performed in GraphPad Prism unless otherwise stated. The statistical details of the experiments are provided in the respective figure legends. Data plot in linear scale were expressed as Mean ± Standard Deviation (SD or median ± interquartile range. Group comparisons were performed using GraphPad Prism version 9.0 and described in each figure legend. Populations were compared using Mann–Whitney U or Wilcoxon Rank-sum test for unpaired comparisons whilst, Student's t-test was applied for paired comparisons, respectively. P values less than or equal 0.05 were considered statistically significant. Kruskal–Wallis and Dunn's post-test were also applied for multiple comparisons in vaccine cohorts. Details pertaining to significance are also provided in the respective figure legends. Spearman R was calculated for the correlation between cytokines and IgG antibodies.

**Suppl. Fig. legends**

**Suppl. Fig. 1 Humoral immune response in 5 different COVID-19 vaccines.** (**a**) Standard curve for 3-plex flow-based assay for Spike S1, RBD and Nucleocapsid antibodies detection from plasma. The FACS plots show the values of different standard points from C0 (low) to C7 (high) for all 3 different antibodies. (**b**) Examples plots from recovered and vaccinated individuals who was positive for SARS-CoV-2 infection and later got COVID-19 vaccine. A clear nucleocapsid antibodies levels can be noticed in infected samples and recovered patient. (**c**) The standard curve for all three antibodies (Spike S1, N and RBD). (**d**) Cellular immune response after first dose of PB mRNA vaccination, significantly reduced levels (p=0.03; Wilcoxon rank-sum test for two samples PrV1 and PoV1 comparisons) of early effector $CD8^+$ T cells after $1^{st}$ dose of PB COVID-19 vaccine. Increased Tregs and $CD56^+$ NK cells after first dose of PB mRNA COVID-19.

**Suppl. Fig. 2 Increased levels of Spike S1 antibody levels in female and ageing declines the antibodies levels of Spike S1 and RBD.** (**a**) Comparison of Spike S1 and RBD levels in male and female. (**b-c**) Age dependent changes in Spike S1 and RBD antibodies levels. Ageing significantly decreased the levels of Spike S1 and RBD levels. Multiple samples comparison was performed using Kruskal Wallis test and post-hoc Dunn's test for multiple comparisons for younger (age group 18-29 Years vs 30-39 Years, 40-49 Years, 50-59 Years and 60-69 Years age group) and with decade increasing age groups. Error bars denote medians and interquartile ranges. *$P < 0.05$; **$P < 0.01$; ***$P < 0.001$; ****$P < 0.0001$.

**Suppl. Fig. 3 scRNA-seq analysis QC for Seurat analysis before data filtration.** All the 28 samples used for the data analysis. (**a-c**) Violin plots before filtering the data show the molecules/cell, genes/cell and % mitochondrial levels in each sample.

**Suppl. Fig. 4 scRNA-seq analysis QC for Seurat analysis after data filtration.** All the 28 samples used for the data analysis. (**a-c**) Violin plots after filtering the data show the molecules/cell, genes/cell and % mitochondrial levels in each sample. (**d**) All the 28 samples used for the PCA guided UMAP analysis and all the 50 PCA were used for UMAP. Samples were batch corrected and Principal component analysis (PCA) of all the analysed samples were shown (Only PC1 and PC2 are shown). (**e**) UMAP analysis for for all the samples used in the study. (**f**) UMAP analysis for PrV1 and PoV2/3 for all the vaccines. (**g**) Manual cell annotation of PBMCs.

**Suppl. Fig. 5 scRNA-seq cell annotation. (a)** Individual cell clusters were manually annotated based on feature bar antibody and (**b**) RNA expression. RNA and protein expression for CD3E, CD4, IL7R, CD8a, MS4A1, CD19, CD79B, IGHD, IGHM, CD27, CD38 and CD14 markers for different T and B cell subsets.

**Suppl. Fig. 6 scRNA-seq cluster confirmation using Azimuth. (a)** Confirmation of $CD4^+$ T cells clusters using azimuth from unsupervised analysis. (**b**) Confirmation of $CD8^+$ T cells clusters using azimuth from unsupervised analysis. (**c**) Confirmation of NKT cells, MAIT and γδ T cells clusters using azimuth from unsupervised analysis. (**d**) Confirmation of NK cells subset clusters using azimuth from unsupervised analysis.

**Suppl. Fig. 7 Comparison of all six vaccines using UMAP analysis.** (**a**) UMAP plots show for Pre-VAC (n=6) and PoV2/3 (n=11) mRNA (BP, MD, and CV), DNA (n=8) (AZ and JJ). (**b-g**) Annotated individual cell clusters are shown in bar diagram for Pre-VAC and post-vaccinated groups for each vaccine types based on material used for inoculation. Major cell clusters such as Tregs, NK, platelets, pMonocytes, CD4+ and CD8+ T cells (total cells) also showed a trend of differentially regulated immune cells for several group.

**Suppl. Fig. 8 Differential gene expression and pathway analysis in DNA and mRNA COVID-19 vaccinations.** (**a**) pMonocytes differential gene expression analysis for mRNA and DNA COVID-19 vaccination (samples were compared with PrV1). Venn diagram that some common gene expression in DNA and mRNA vaccination Metascape tool was used performed gene enrichment analysis for different GO pathways to understand the function of genes upregulated in pMonocytes. (**b**) NKIII cells differential gene expression analysis for mRNA and DNA COVID-19 vaccination (samples were compared with PrV1) and gene pathway analysis. (**b**) NKT cells differential gene expression analysis for mRNA and DNA COVID-19 vaccination (samples were compared with PrV1) and gene pathway analysis.

**Suppl. Fig. 9 Differential gene expression analysis of pMonocytes.** (**a**) Using Metascape tool, we performed gene enrichment analysis for different GO pathways to understand the function of genes upregulated in pMonocytes specific cell cluster. In pMonocytes, several pathways were regulated with defense response to fungus and leukocyte cell-cell adhesion. (**b-g**) Overall differential gene expression in different cell clusters in individual vaccines before and after vaccination. (a) AC007952, S100AB, IER3, IL1B, CD83 and RGCC in AZ vaccinated individuals. (b) S100AB, IER3, IL1B, VIM, AL138963.4 and NFKBIA in CV vaccinated individuals. (c) HSPAS, SRGN, PLAUR, RGCC, CXCL2 and PHLDA1 in MD vaccinated individuals. **Differential gene expression analysis before and after AZ vaccination.** (**a**) AC007952.4 and S100AB genes were upregulated, whilst IER3, IL1B, CD83, RGCC were significantly downregulated after AZ vaccination in cMonocytes. (**b**) RASGEF1B, TAGAP, FOSB, CD83, PLAUR, IER3 were significantly downregulated after AZ vaccination in pMonocytes. (**c**) JUN, H3F3B, GADD45B and IER2genes were upregulated, whilst TSC22D3 and TNFAIP3 were significantly downregulated after AZ vaccination in γδ T cells.

**Suppl. Fig. 10 RNA-seq analysis pipeline by "SeqGeq", identification of specific T cell subsets and protein marker expression for T cell subsets.** (**a**) All the 26 samples from cell ranger output in an expression matrix files (.h5 – HDF5 common output file from 10x Genomics platform; "filtered_gene_bc_matrices.h5") were concatenated to analyse single cell RNA-sequencing (scRNA-Seq). First individual samples were normalized transcript per 10,000 to adjust the data to a common scale then all the samples were concatenated.  The later samples were assessed for quality control (QC) to remove the ambiguous or questionable data points. Three main steps: (i) visualizing, (ii) assessing, and (iii) filtering was performed in this order at an early stage of the analysis process. The first one in gene view and the last two in cell view. First, we checked how the overall cells look in all the samples. The first QC plot was used remove outlier events (empty wells or doublets): x-axis library size vs y-axis gene expressed (both in log scale). The second plot (x-axis total reads vs y-axis cells expressing) was to remove genes expressed by few cells or all cells (both on log scale). The third QC plot (x-axis cells expressing vs y-axis dispersion) was used to remove genes with low variance in expression levels. Dispersion index represents variance divided by mean. Then using Seurat plugin, dispersed genes were used for dimensional reduction plots for unbiased cell clustering using PCA guided UMAP analysis. (**b**) UMAP plots show different $CD4^+$ T cell subsets (left) and UMAP plots show different $CD8^+$ T cell subsets (right). (**c**) UMAP plots show individual cell specific marker protein and RNA levels to identify the identity of each cell cluster. CD3D, CD4, CD8, CD14, CD16a, CD11c, CD19 and CD56 or NCAM1 for T cells, monocytes, dendritic cells, and NK cells. UMAP plots show different naïve or memory marker of either $CD4^+$ or $CD8^+$ T cells on RNA and proteins such as CCR7, PTPRC, SELL, CD27, LEF1, ITGA6 and PRDM1. (**d**) UMAP plots shows the distribution of different vaccines. (**e**) Differential expression of genes at mRNA and protein levels in PrV1 vs PoV2-AZ. Significant increased levels of CD2 & ITGB2 levels (Benjamini-Hochberg multiple correction test and false discovery rate test (FDR) value $q<0.05$).

**Suppl. Fig. 11 Expression of activation markers in DNA and RNA vaccination.** (**a**) UMAP analysis of individual markers (CD2, CD244, CD58, CD48, ITGB2, CCR5 and MALAT1) at protein levels. (**b**) Violin plots represent the expression of CD2, CD244, CD58, CD48, CCR5 and ITGB2 in individual cell clusters.

**Suppl. Fig. 12 Immunophenotyping of Pre-VAC and PoV2/PoV3 COVID-19 vaccinated individuals based on 14 colour TBNK and monocytes flow cytometry panel.** (**a**) UMAP analysis show the expression of individual antibody abundance in PBMCs for unvaccinated and vaccinated individuals after different vaccination. (**b**) Expression of CD4 and CD8 ration in PrV1, PoV2 and PoV3. (**b**) Expression of CCR7 and CD45RA to assess the naïve memory T cell marker for $CD4^+$ and $CD8^+$ T cells in Pre-VAC, PoV3 and PoV3.

**Suppl. Fig. 13 Antibodies and cytokine data correlation with age and sex.** (**a**) Detection of 13-different cytokines. Data represented in colour code information for individual vaccination. (**b-c**) Detection of 13-different cytokines. Data represented in colour code information for individual vaccination. (**d**) Correlation analysis of cytokines in PoV3 vaccinated individuals (PoV2: 3 - 6 months).

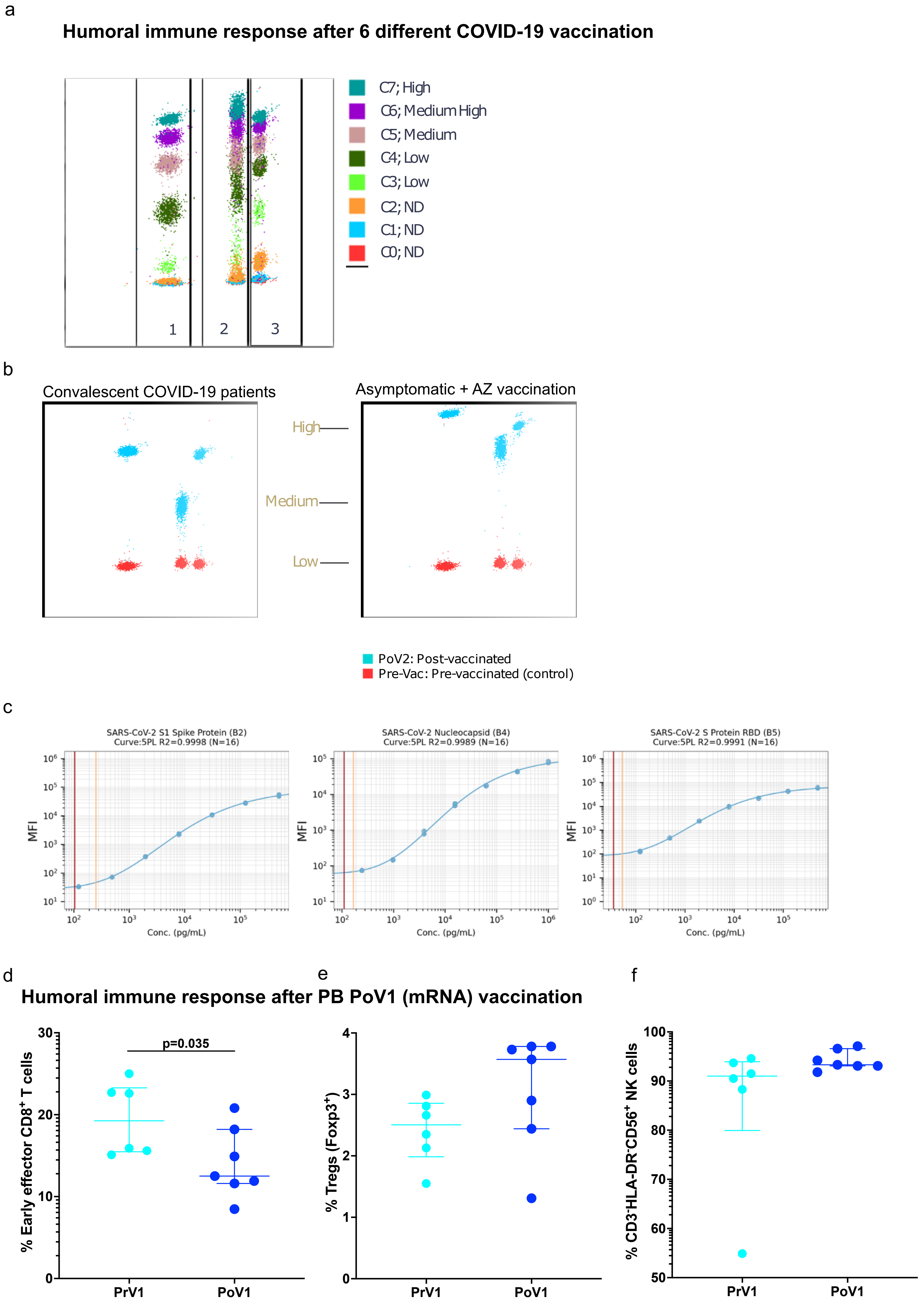
a
Humoral immune response after 6 different COVID-19 vaccination
1
2
3
C7; High
C6; Medium High
C5; Medium
C4; Low
C3; Low
C2; ND
C1; ND
C0; ND
b
Convalescent COVID-19 patients
Asymptomatic + AZ vaccination
High
Medium
Low
PoV2: Post-vaccinated
Pre-Vac: Pre-vaccinated (control)
c
SARS-CoV-2 S1 Spike Protein (B2)
Curve:5PL R2=0.9998 (N=16)
SARS-CoV-2 Nucleocapsid (B4)
Curve:5PL R2=0.9989 (N=16)
SARS-CoV-2 S Protein RBD (B5)
Curve:5PL R2=0.9991 (N=16)
MFI
Conc. (pg/mL)
d
e
f
Humoral immune response after PB PoV1 (mRNA) vaccination
% Early effector CD8+ T cells
p=0.035
% Tregs (Foxp3+)
% CD3-HLA-DR-CD56+ NK cells
PrV1
PoV1

Suppl. Fig. 1

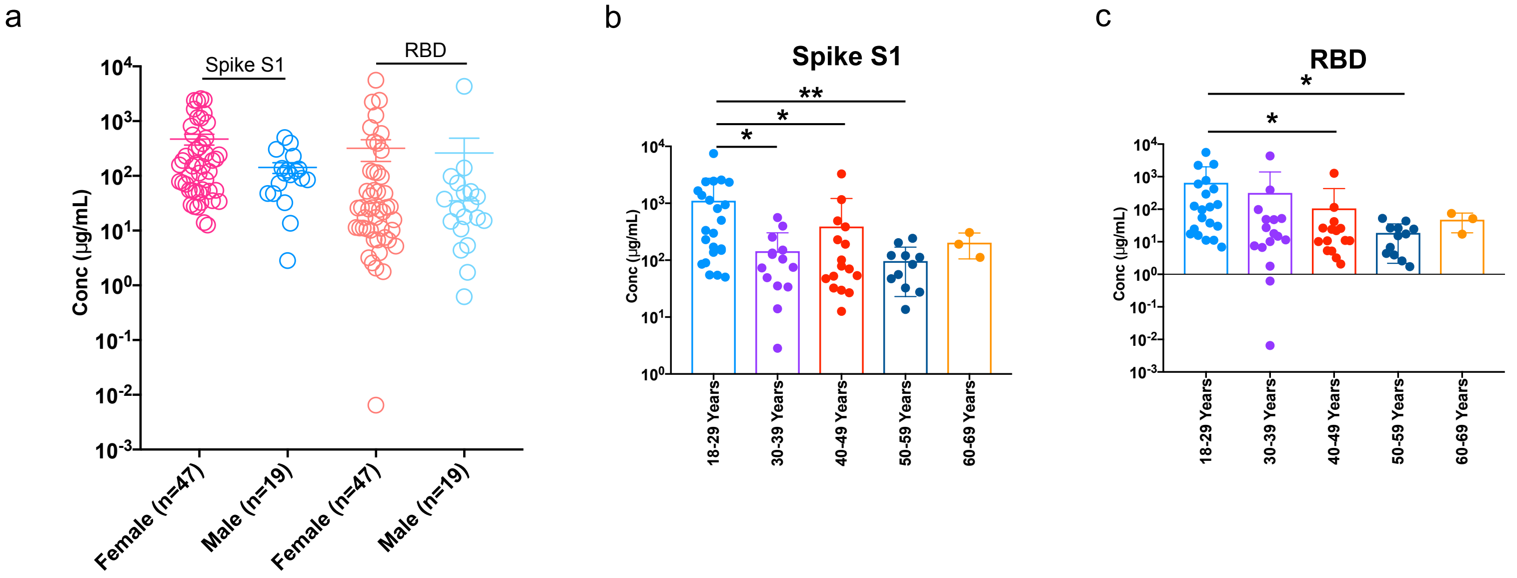
a
Spike S1
RBD
Conc (µg/mL)
Female (n=47)
Male (n=19)
Female (n=47)
Male (n=19)
b
Spike S1
Conc (µg/mL)
18-29 Years
30-39 Years
40-49 Years
50-59 Years
60-69 Years
c
RBD
Conc (µg/mL)
18-29 Years
30-39 Years
40-49 Years
50-59 Years
60-69 Years

Suppl. Fig. 2

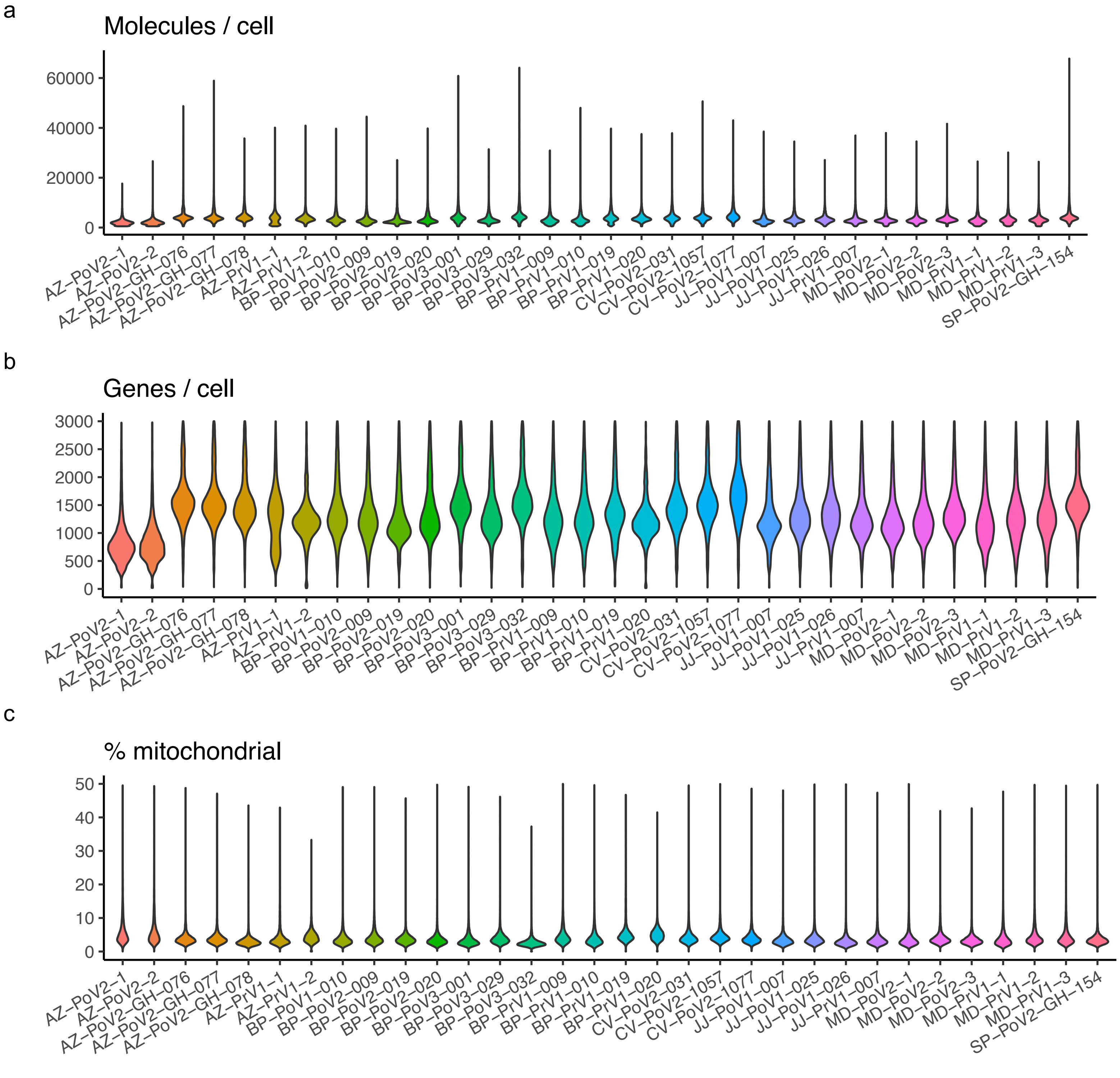
a
Molecules / cell
60000
40000
20000
0
AZ–PoV2–1
AZ–PoV2–2
AZ–PoV2–GH–076
AZ–PoV2–GH–077
AZ–PoV2–GH–078
AZ–PrV1–1
AZ–PrV1–2
BP–PoV1–010
BP–PoV2–009
BP–PoV2–019
BP–PoV2–020
BP–PoV3–001
BP–PoV3–029
BP–PoV3–032
BP–PrV1–009
BP–PrV1–010
BP–PrV1–019
BP–PrV1–020
CV–PoV2–031
CV–PoV2–1057
CV–PoV2–1077
JJ–PoV1–007
JJ–PoV1–025
JJ–PoV1–026
JJ–PrV1–007
MD–PoV2–1
MD–PoV2–2
MD–PoV2–3
MD–PrV1–1
MD–PrV1–2
MD–PrV1–3
SP–PoV2–GH–154
b
Genes / cell
3000
2500
2000
1500
1000
500
0
AZ–PoV2–1
AZ–PoV2–2
AZ–PoV2–GH–076
AZ–PoV2–GH–077
AZ–PoV2–GH–078
AZ–PrV1–1
AZ–PrV1–2
BP–PoV1–010
BP–PoV2–009
BP–PoV2–019
BP–PoV2–020
BP–PoV3–001
BP–PoV3–029
BP–PoV3–032
BP–PrV1–009
BP–PrV1–010
BP–PrV1–019
BP–PrV1–020
CV–PoV2–031
CV–PoV2–1057
CV–PoV2–1077
JJ–PoV1–007
JJ–PoV1–025
JJ–PoV1–026
JJ–PrV1–007
MD–PoV2–1
MD–PoV2–2
MD–PoV2–3
MD–PrV1–1
MD–PrV1–2
MD–PrV1–3
SP–PoV2–GH–154
c
% mitochondrial
50
40
30
20
10
0
AZ–PoV2–1
AZ–PoV2–2
AZ–PoV2–GH–076
AZ–PoV2–GH–077
AZ–PoV2–GH–078
AZ–PrV1–1
AZ–PrV1–2
BP–PoV1–010
BP–PoV2–009
BP–PoV2–019
BP–PoV2–020
BP–PoV3–001
BP–PoV3–029
BP–PoV3–032
BP–PrV1–009
BP–PrV1–010
BP–PrV1–019
BP–PrV1–020
CV–PoV2–031
CV–PoV2–1057
CV–PoV2–1077
JJ–PoV1–007
JJ–PoV1–025
JJ–PoV1–026
JJ–PrV1–007
MD–PoV2–1
MD–PoV2–2
MD–PoV2–3
MD–PrV1–1
MD–PrV1–2
MD–PrV1–3
SP–PoV2–GH–154

Suppl. Fig. 3

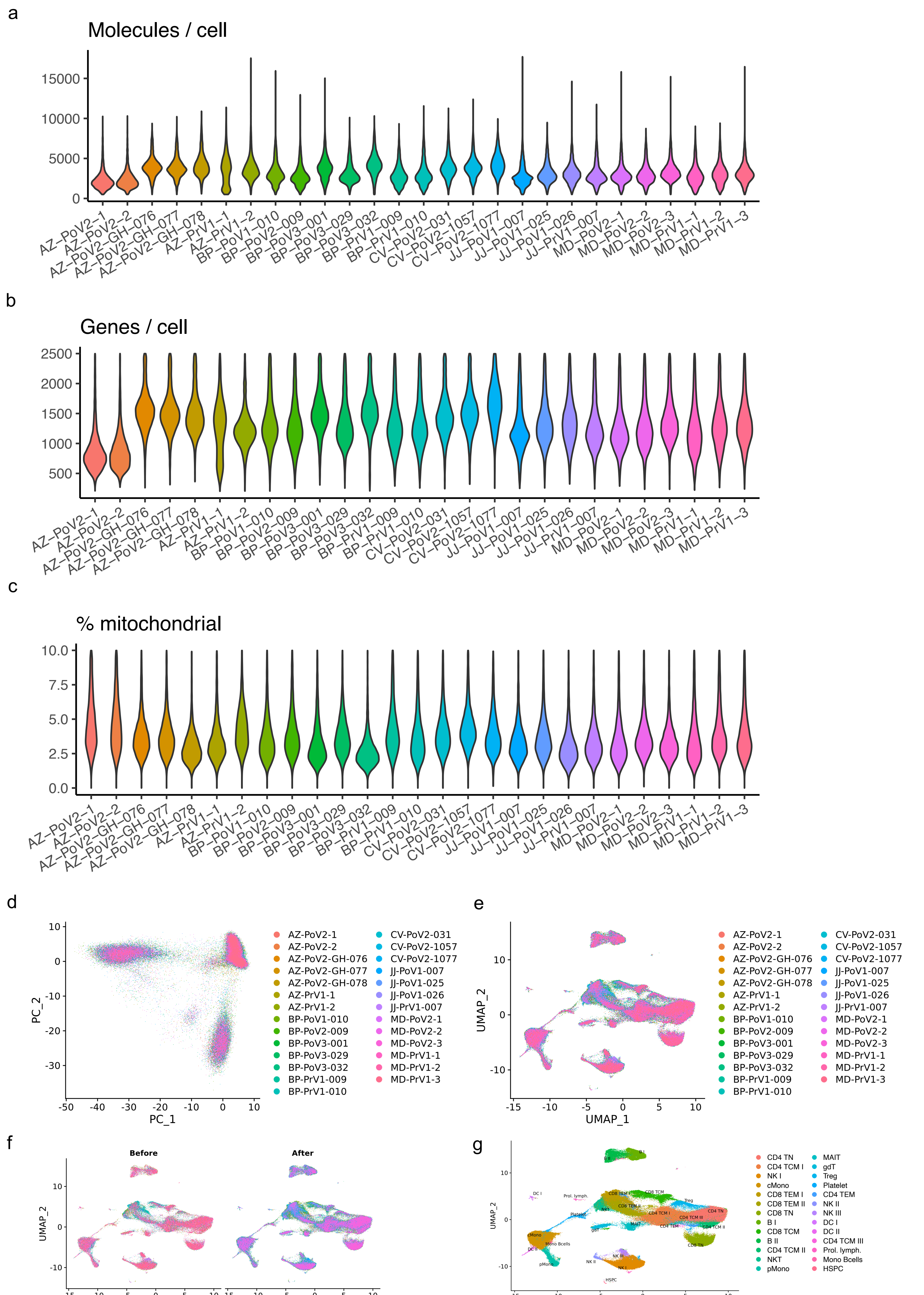
a
Molecules / cell
0
5000
10000
15000
AZ-PoV2-1 AZ-PoV2-2 AZ-PoV2-GH-076 AZ-PoV2-GH-077 AZ-PoV2-GH-078 AZ-PrV1-1 AZ-PrV1-2 BP-PoV1-010 BP-PoV2-009 BP-PoV3-001 BP-PoV3-029 BP-PoV3-032 BP-PrV1-009 BP-PrV1-010 CV-PoV2-031 CV-PoV2-1057 CV-PoV2-1077 JJ-PoV1-007 JJ-PoV1-025 JJ-PoV1-026 JJ-PrV1-007 MD-PoV2-1 MD-PoV2-2 MD-PoV2-3 MD-PrV1-1 MD-PrV1-2 MD-PrV1-3
b
Genes / cell
500
1000
1500
2000
2500
c
% mitochondrial
0.0
2.5
5.0
7.5
10.0
d
PC_1
PC_2
e
UMAP_1
UMAP_2
AZ-PoV2-1
AZ-PoV2-2
AZ-PoV2-GH-076
AZ-PoV2-GH-077
AZ-PoV2-GH-078
AZ-PrV1-1
AZ-PrV1-2
BP-PoV1-010
BP-PoV2-009
BP-PoV3-001
BP-PoV3-029
BP-PoV3-032
BP-PrV1-009
BP-PrV1-010
CV-PoV2-031
CV-PoV2-1057
CV-PoV2-1077
JJ-PoV1-007
JJ-PoV1-025
JJ-PoV1-026
JJ-PrV1-007
MD-PoV2-1
MD-PoV2-2
MD-PoV2-3
MD-PrV1-1
MD-PrV1-2
MD-PrV1-3
f
Before
After
UMAP_1
UMAP_2
g
UMAP_1
UMAP_2
CD4 TN
CD4 TCM I
NK I
cMono
CD8 TEM I
CD8 TEM II
CD8 TN
B I
CD8 TCM
B II
CD4 TCM II
NKT
pMono
MAIT
gdT
Treg
Platelet
CD4 TEM
NK II
NK III
DC I
DC II
CD4 TCM III
Prol. lymph.
Mono Bcells
HSPC

Suppl. Fig. 4

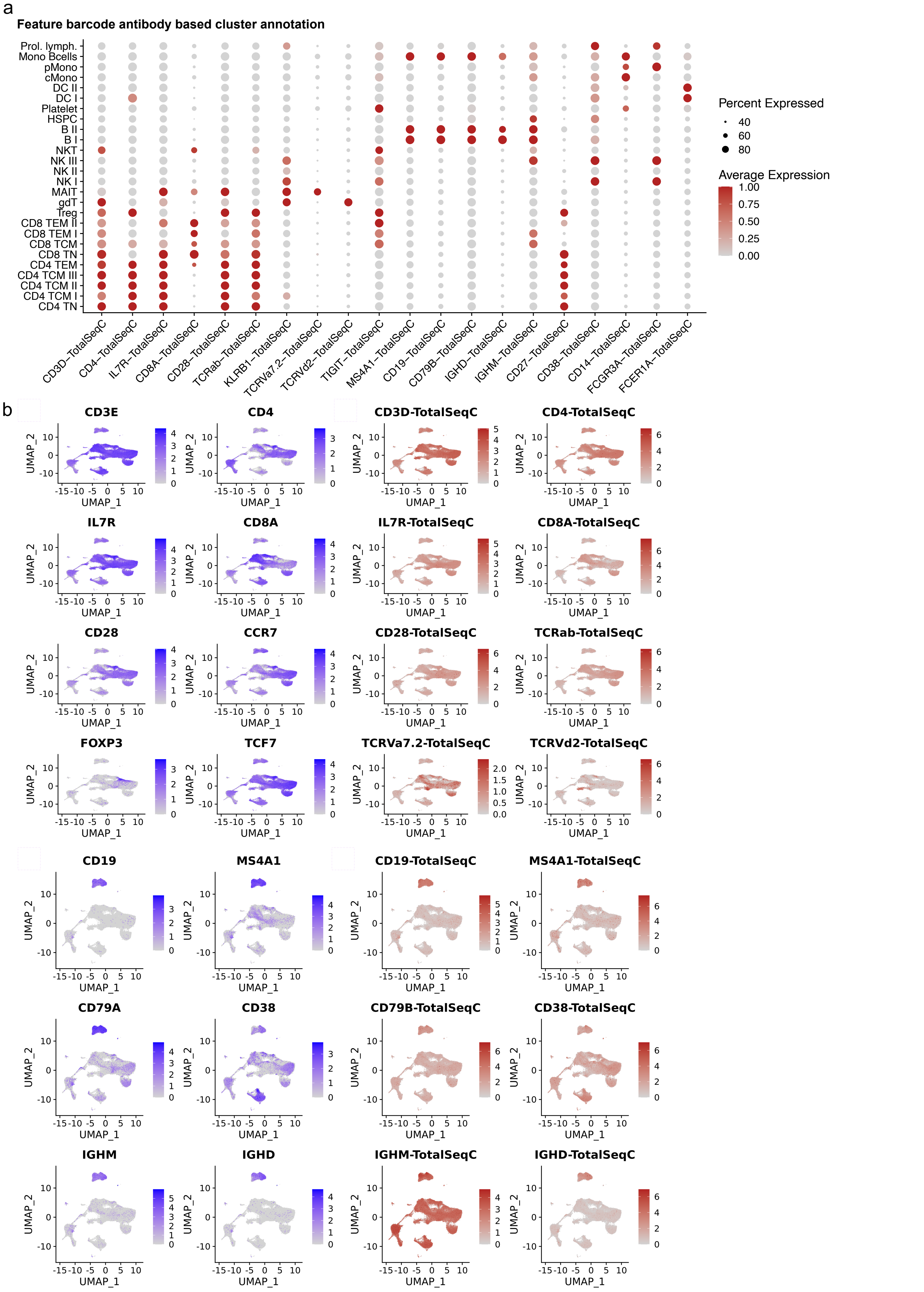
a
Feature barcode antibody based cluster annotation
Prol. lymph.
Mono Bcells
pMono
cMono
DC II
DC I
Platelet
HSPC
B II
B I
NKT
NK III
NK II
NK I
MAIT
gdT
Treg
CD8 TEM II
CD8 TEM I
CD8 TCM
CD8 TN
CD4 TEM
CD4 TCM III
CD4 TCM II
CD4 TCM I
CD4 TN
CD3D–TotalSeqC
CD4–TotalSeqC
IL7R–TotalSeqC
CD8A–TotalSeqC
CD28–TotalSeqC
TCRab–TotalSeqC
KLRB1–TotalSeqC
TCRVa7.2–TotalSeqC
TCRVd2–TotalSeqC
TIGIT–TotalSeqC
MS4A1–TotalSeqC
CD19–TotalSeqC
CD79B–TotalSeqC
IGHD–TotalSeqC
IGHM–TotalSeqC
CD27–TotalSeqC
CD38–TotalSeqC
CD14–TotalSeqC
FCGR3A–TotalSeqC
FCER1A–TotalSeqC
Percent Expressed
40
60
80
Average Expression
1.00
0.75
0.50
0.25
0.00
b
CD3E
CD4
CD3D-TotalSeqC
CD4-TotalSeqC
IL7R
CD8A
IL7R-TotalSeqC
CD8A-TotalSeqC
CD28
CCR7
CD28-TotalSeqC
TCRab-TotalSeqC
FOXP3
TCF7
TCRVa7.2-TotalSeqC
TCRVd2-TotalSeqC
CD19
MS4A1
CD19-TotalSeqC
MS4A1-TotalSeqC
CD79A
CD38
CD79B-TotalSeqC
CD38-TotalSeqC
IGHM
IGHD
IGHM-TotalSeqC
IGHD-TotalSeqC
UMAP_1
UMAP_2

Suppl. Fig. 5

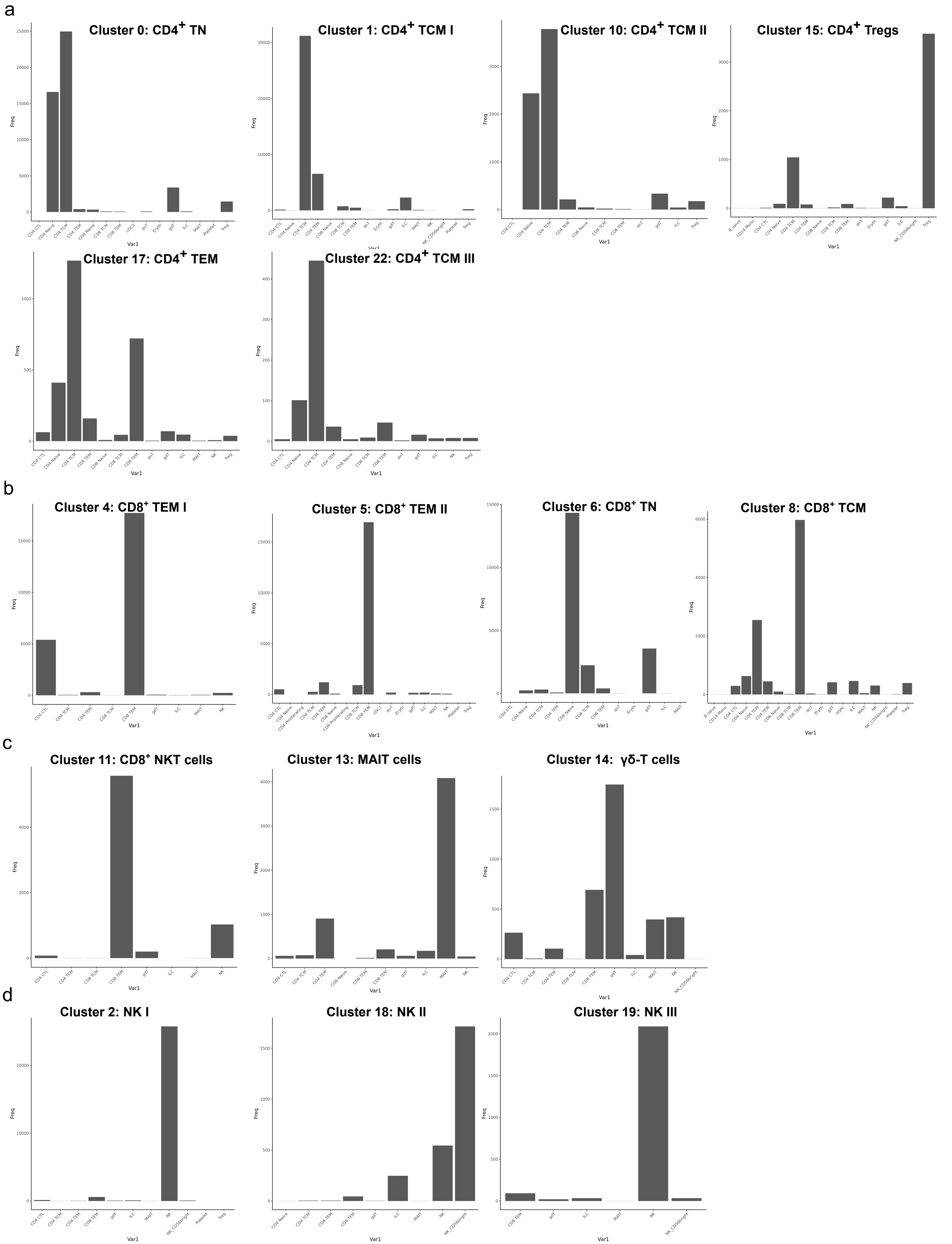
a
Cluster 0: CD4+ TN
Cluster 1: CD4+ TCM I
Cluster 10: CD4+ TCM II
Cluster 15: CD4+ Tregs
Cluster 17: CD4+ TEM
Cluster 22: CD4+ TCM III
b
Cluster 4: CD8+ TEM I
Cluster 5: CD8+ TEM II
Cluster 6: CD8+ TN
Cluster 8: CD8+ TCM
c
Cluster 11: CD8+ NKT cells
Cluster 13: MAIT cells
Cluster 14: γδ-T cells
d
Cluster 2: NK I
Cluster 18: NK II
Cluster 19: NK III
Freq
Var1

Suppl. Fig. 6

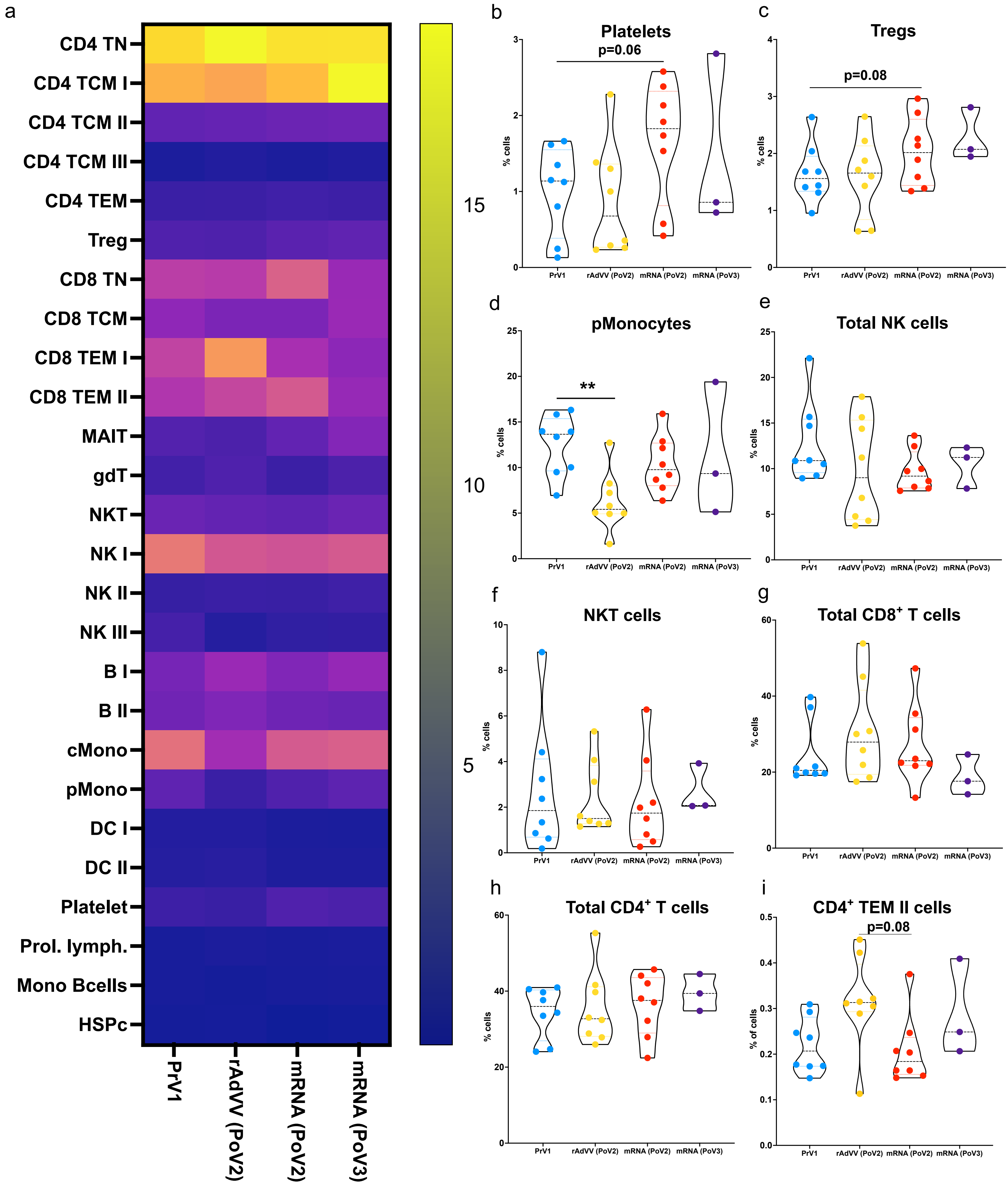
a
CD4 TN
CD4 TCM I
CD4 TCM II
CD4 TCM III
CD4 TEM
Treg
CD8 TN
CD8 TCM
CD8 TEM I
CD8 TEM II
MAIT
gdT
NKT
NK I
NK II
NK III
B I
B II
cMono
pMono
DC I
DC II
Platelet
Prol. lymph.
Mono Bcells
HSPc
PrV1
rAdVV (PoV2)
mRNA (PoV2)
mRNA (PoV3)
15
10
5
b
Platelets
p=0.06
c
Tregs
p=0.08
d
pMonocytes
**
e
Total NK cells
f
NKT cells
g
Total CD8+ T cells
h
Total CD4+ T cells
i
CD4+ TEM II cells
p=0.08
% cells
% of cells

Suppl. Fig. 7

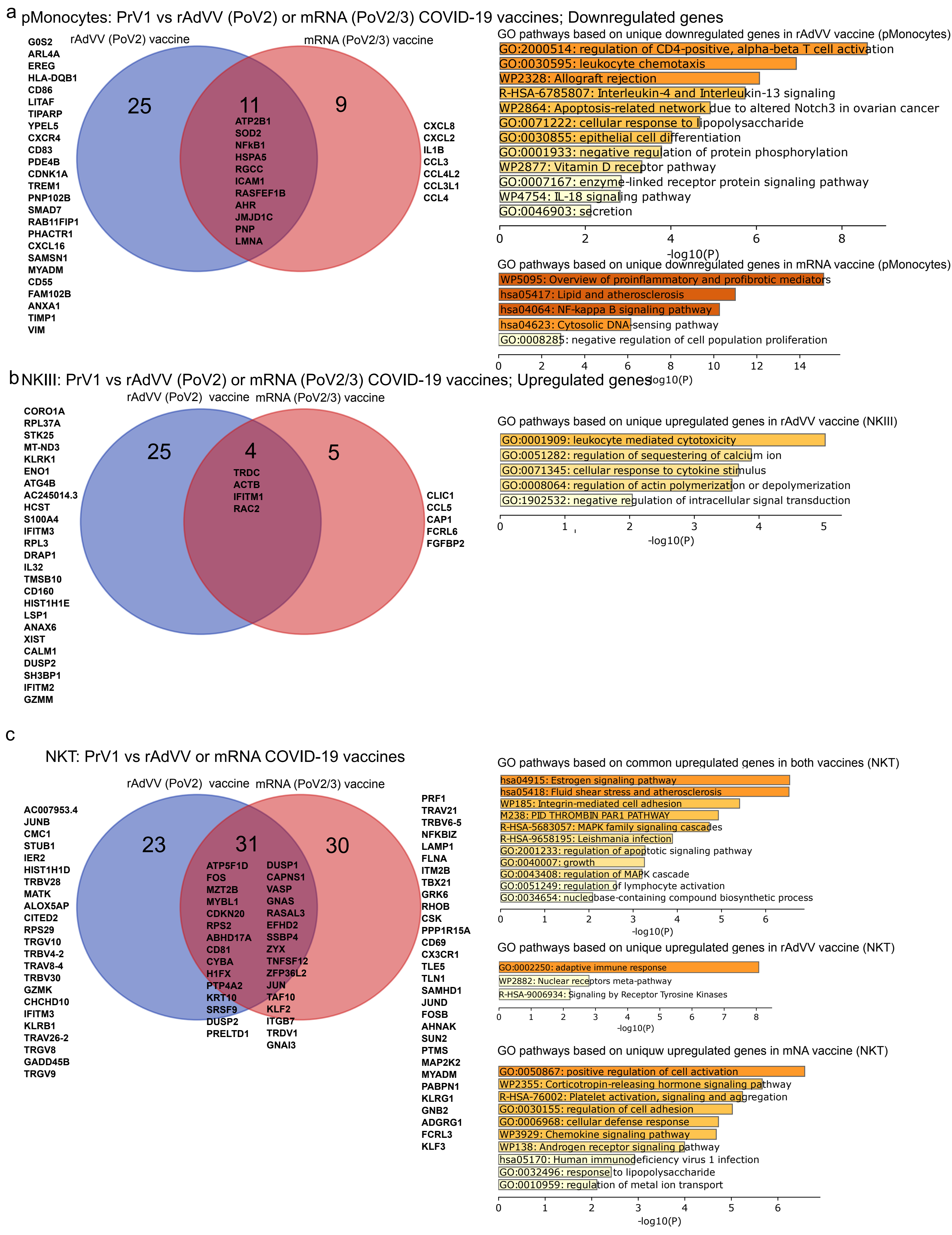
a pMonocytes: PrV1 vs rAdVV (PoV2) or mRNA (PoV2/3) COVID-19 vaccines; Downregulated genes
rAdVV (PoV2) vaccine
mRNA (PoV2/3) vaccine
25
11
9
G0S2 ARL4A EREG HLA-DQB1 CD86 LITAF TIPARP YPEL5 CXCR4 CD83 PDE4B CDNK1A TREM1 PNP102B SMAD7 RAB11FIP1 PHACTR1 CXCL16 SAMSN1 MYADM CD55 FAM102B ANXA1 TIMP1 VIM
ATP2B1 SOD2 NFkB1 HSPA5 RGCC ICAM1 RASFEF1B AHR JMJD1C PNP LMNA
CXCL8 CXCL2 IL1B CCL3 CCL4L2 CCL3L1 CCL4
GO pathways based on unique downregulated genes in rAdVV vaccine (pMonocytes)
GO:2000514: regulation of CD4-positive, alpha-beta T cell activation
GO:0030595: leukocyte chemotaxis
WP2328: Allograft rejection
R-HSA-6785807: Interleukin-4 and Interleukin-13 signaling
WP2864: Apoptosis-related network due to altered Notch3 in ovarian cancer
GO:0071222: cellular response to lipopolysaccharide
GO:0030855: epithelial cell differentiation
GO:0001933: negative regulation of protein phosphorylation
WP2877: Vitamin D receptor pathway
GO:0007167: enzyme-linked receptor protein signaling pathway
WP4754: IL-18 signaling pathway
GO:0046903: secretion
0 2 4 6 8
-log10(P)
GO pathways based on unique downregulated genes in mRNA vaccine (pMonocytes)
WP5095: Overview of proinflammatory and profibrotic mediators
hsa05417: Lipid and atherosclerosis
hsa04064: NF-kappa B signaling pathway
hsa04623: Cytosolic DNA-sensing pathway
GO:0008285: negative regulation of cell population proliferation
0 2 4 6 8 10 12 14
-log10(P)
b NKIII: PrV1 vs rAdVV (PoV2) or mRNA (PoV2/3) COVID-19 vaccines; Upregulated genes
rAdVV (PoV2) vaccine
mRNA (PoV2/3) vaccine
25
4
5
CORO1A RPL37A STK25 MT-ND3 KLRK1 ENO1 ATG4B AC245014.3 HCST S100A4 IFITM3 RPL3 DRAP1 IL32 TMSB10 CD160 HIST1H1E LSP1 ANAX6 XIST CALM1 DUSP2 SH3BP1 IFITM2 GZMM
TRDC ACTB IFITM1 RAC2
CLIC1 CCL5 CAP1 FCRL6 FGFBP2
GO pathways based on unique upregulated genes in rAdVV vaccine (NKIII)
GO:0001909: leukocyte mediated cytotoxicity
GO:0051282: regulation of sequestering of calcium ion
GO:0071345: cellular response to cytokine stimulus
GO:0008064: regulation of actin polymerization or depolymerization
GO:1902532: negative regulation of intracellular signal transduction
0 1 2 3 4 5
-log10(P)
c
NKT: PrV1 vs rAdVV or mRNA COVID-19 vaccines
rAdVV (PoV2) vaccine
mRNA (PoV2/3) vaccine
23
31
30
AC007953.4 JUNB CMC1 STUB1 IER2 HIST1H1D TRBV28 MATK ALOX5AP CITED2 RPS29 TRGV10 TRBV4-2 TRAV8-4 TRBV30 GZMK CHCHD10 IFITM3 KLRB1 TRAV26-2 TRGV8 GADD45B TRGV9
ATP5F1D FOS MZT2B MYBL1 CDKN20 RPS2 ABHD17A CD81 CYBA H1FX PTP4A2 KRT10 SRSF9 DUSP2 PRELTD1
DUSP1 CAPNS1 VASP GNAS RASAL3 EFHD2 SSBP4 ZYX TNFSF12 ZFP36L2 JUN TAF10 KLF2 ITGB7 TRDV1 GNAI3
PRF1 TRAV21 TRBV6-5 NFKBIZ LAMP1 FLNA ITM2B TBX21 GRK6 RHOB CSK PPP1R15A CD69 CX3CR1 TLE5 TLN1 SAMHD1 JUND FOSB AHNAK SUN2 PTMS MAP2K2 MYADM PABPN1 KLRG1 GNB2 ADGRG1 FCRL3 KLF3
GO pathways based on common upregulated genes in both vaccines (NKT)
hsa04915: Estrogen signaling pathway
hsa05418: Fluid shear stress and atherosclerosis
WP185: Integrin-mediated cell adhesion
M238: PID THROMBIN PAR1 PATHWAY
R-HSA-5683057: MAPK family signaling cascades
R-HSA-9658195: Leishmania infection
GO:2001233: regulation of apoptotic signaling pathway
GO:0040007: growth
GO:0043408: regulation of MAPK cascade
GO:0051249: regulation of lymphocyte activation
GO:0034654: nucleobase-containing compound biosynthetic process
0 1 2 3 4 5 6
-log10(P)
GO pathways based on unique upregulated genes in rAdVV vaccine (NKT)
GO:0002250: adaptive immune response
WP2882: Nuclear receptors meta-pathway
R-HSA-9006934: Signaling by Receptor Tyrosine Kinases
0 1 2 3 4 5 6 7 8
-log10(P)
GO pathways based on uniquw upregulated genes in mNA vaccine (NKT)
GO:0050867: positive regulation of cell activation
WP2355: Corticotropin-releasing hormone signaling pathway
R-HSA-76002: Platelet activation, signaling and aggregation
GO:0030155: regulation of cell adhesion
GO:0006968: cellular defense response
WP3929: Chemokine signaling pathway
WP138: Androgen receptor signaling pathway
hsa05170: Human immunodeficiency virus 1 infection
GO:0032496: response to lipopolysaccharide
GO:0010959: regulation of metal ion transport
0 1 2 3 4 5 6
-log10(P)

Suppl. Fig. 8

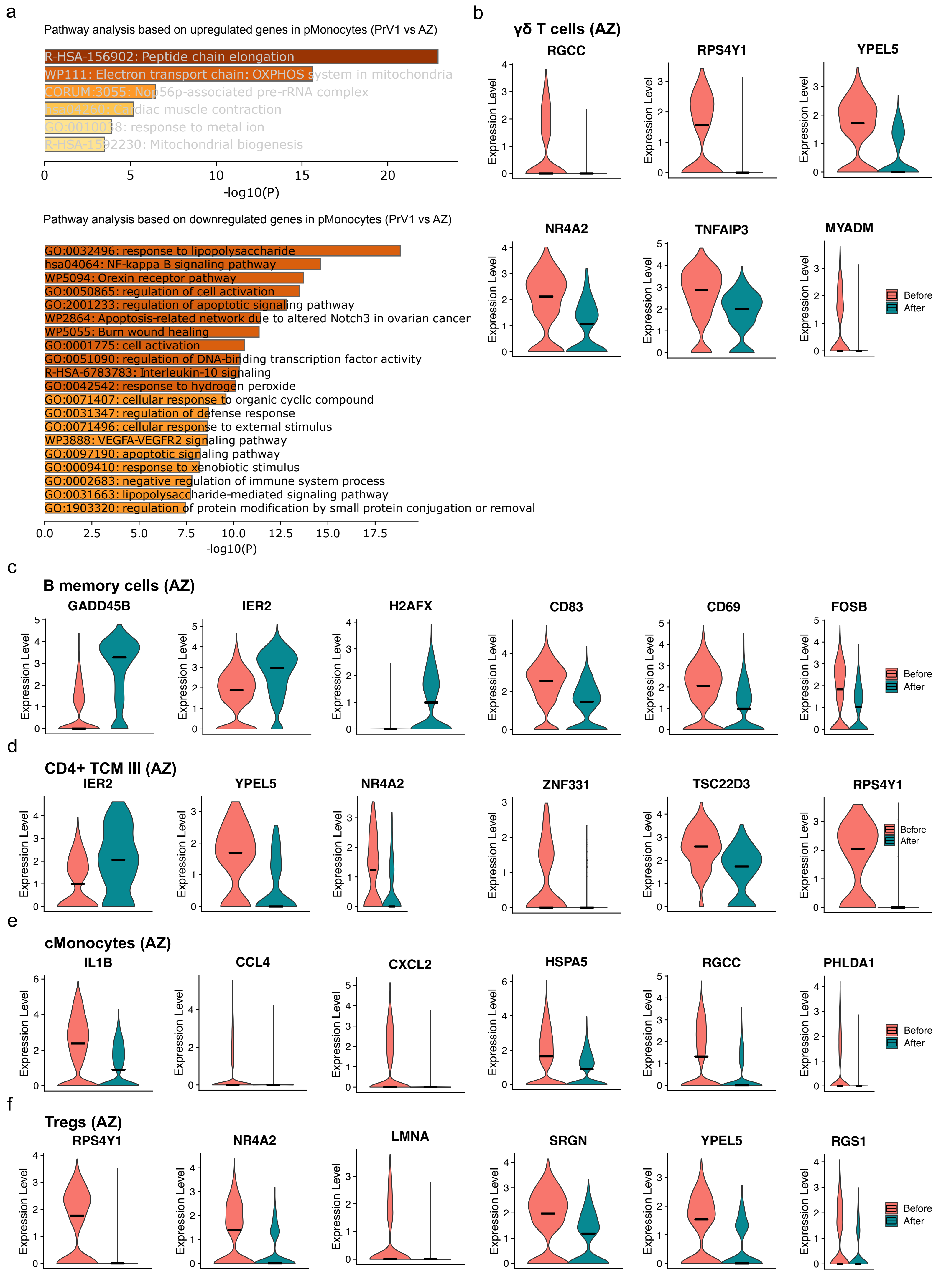
a
Pathway analysis based on upregulated genes in pMonocytes (PrV1 vs AZ)
R-HSA-156902: Peptide chain elongation
WP111: Electron transport chain: OXPHOS system in mitochondria
CORUM:3055: Nop56p-associated pre-rRNA complex
hsa04260: Cardiac muscle contraction
GO:0010038: response to metal ion
R-HSA-1592230: Mitochondrial biogenesis
-log10(P)
Pathway analysis based on downregulated genes in pMonocytes (PrV1 vs AZ)
GO:0032496: response to lipopolysaccharide
hsa04064: NF-kappa B signaling pathway
WP5094: Orexin receptor pathway
GO:0050865: regulation of cell activation
GO:2001233: regulation of apoptotic signaling pathway
WP2864: Apoptosis-related network due to altered Notch3 in ovarian cancer
WP5055: Burn wound healing
GO:0001775: cell activation
GO:0051090: regulation of DNA-binding transcription factor activity
R-HSA-6783783: Interleukin-10 signaling
GO:0042542: response to hydrogen peroxide
GO:0071407: cellular response to organic cyclic compound
GO:0031347: regulation of defense response
GO:0071496: cellular response to external stimulus
WP3888: VEGFA-VEGFR2 signaling pathway
GO:0097190: apoptotic signaling pathway
GO:0009410: response to xenobiotic stimulus
GO:0002683: negative regulation of immune system process
GO:0031663: lipopolysaccharide-mediated signaling pathway
GO:1903320: regulation of protein modification by small protein conjugation or removal
-log10(P)
b
γδ T cells (AZ)
RGCC
RPS4Y1
YPEL5
NR4A2
TNFAIP3
MYADM
Expression Level
Before
After
c
B memory cells (AZ)
GADD45B
IER2
H2AFX
CD83
CD69
FOSB
d
CD4+ TCM III (AZ)
IER2
YPEL5
NR4A2
ZNF331
TSC22D3
RPS4Y1
e
cMonocytes (AZ)
IL1B
CCL4
CXCL2
HSPA5
RGCC
PHLDA1
f
Tregs (AZ)
RPS4Y1
NR4A2
LMNA
SRGN
YPEL5
RGS1

Suppl. Fig. 9

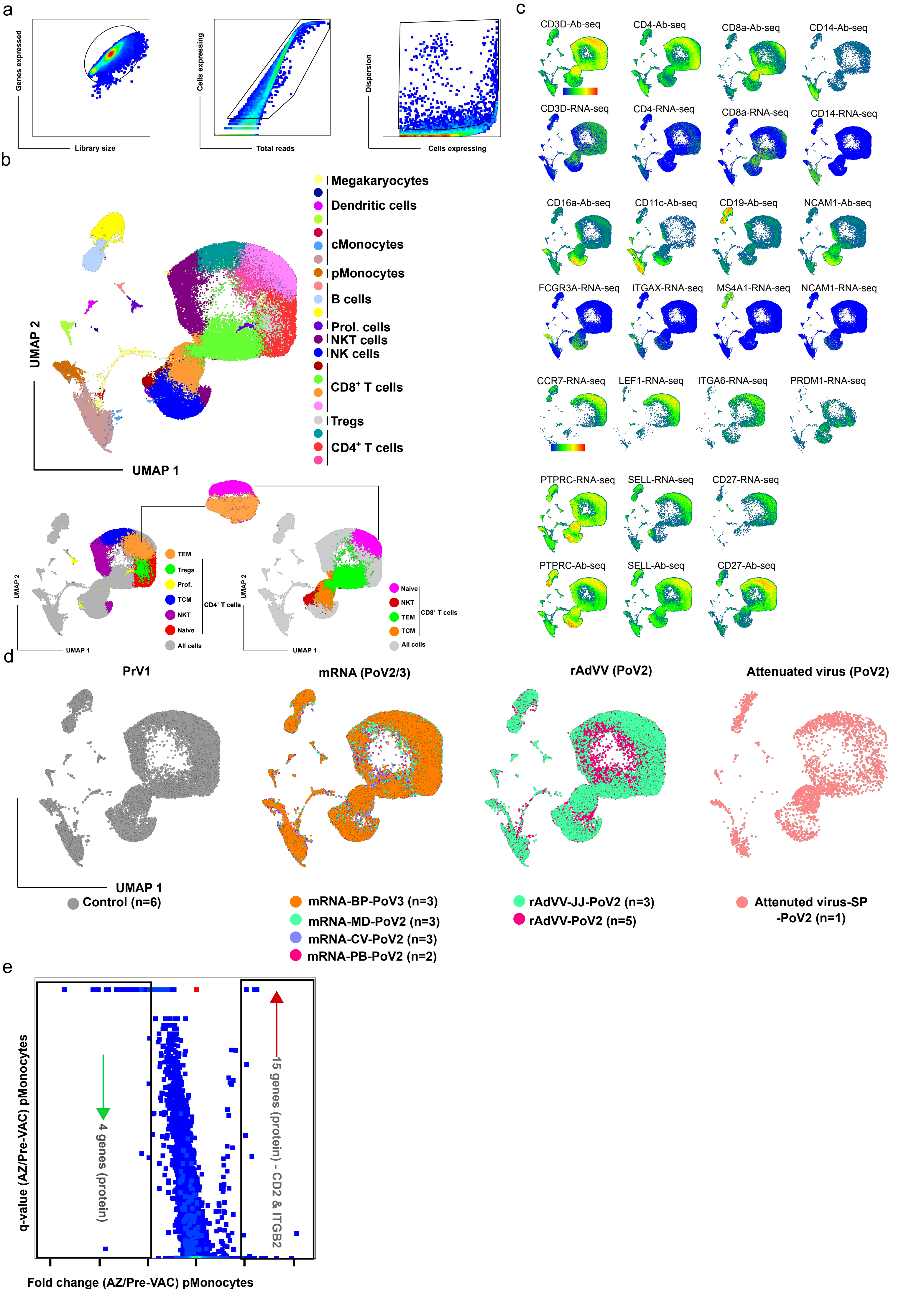
a
Genes expressed
Library size
Cells expressing
Total reads
Dispersion
Cells expressing
b
UMAP 2
UMAP 1
Megakaryocytes
Dendritic cells
cMonocytes
pMonocytes
B cells
Prol. cells
NKT cells
NK cells
CD8⁺ T cells
Tregs
CD4⁺ T cells
TEM
Tregs
Prof.
TCM
NKT
Naive
All cells
CD4⁺ T cells
Naive
NKT
TEM
TCM
All cells
CD8⁺ T cells
c
CD3D-Ab-seq
CD4-Ab-seq
CD8a-Ab-seq
CD14-Ab-seq
CD3D-RNA-seq
CD4-RNA-seq
CD8a-RNA-seq
CD14-RNA-seq
CD16a-Ab-seq
CD11c-Ab-seq
CD19-Ab-seq
NCAM1-Ab-seq
FCGR3A-RNA-seq
ITGAX-RNA-seq
MS4A1-RNA-seq
NCAM1-RNA-seq
CCR7-RNA-seq
LEF1-RNA-seq
ITGA6-RNA-seq
PRDM1-RNA-seq
PTPRC-RNA-seq
SELL-RNA-seq
CD27-RNA-seq
PTPRC-Ab-seq
SELL-Ab-seq
CD27-Ab-seq
d
PrV1
mRNA (PoV2/3)
rAdVV (PoV2)
Attenuated virus (PoV2)
UMAP 1
Control (n=6)
mRNA-BP-PoV3 (n=3)
mRNA-MD-PoV2 (n=3)
mRNA-CV-PoV2 (n=3)
mRNA-PB-PoV2 (n=2)
rAdVV-JJ-PoV2 (n=3)
rAdVV-PoV2 (n=5)
Attenuted virus-SP
-PoV2 (n=1)
e
q-value (AZ/Pre-VAC) pMonocytes
4 genes (protein)
15 genes (protein) - CD2 & ITGB2
Fold change (AZ/Pre-VAC) pMonocytes

Suppl. Fig. 10

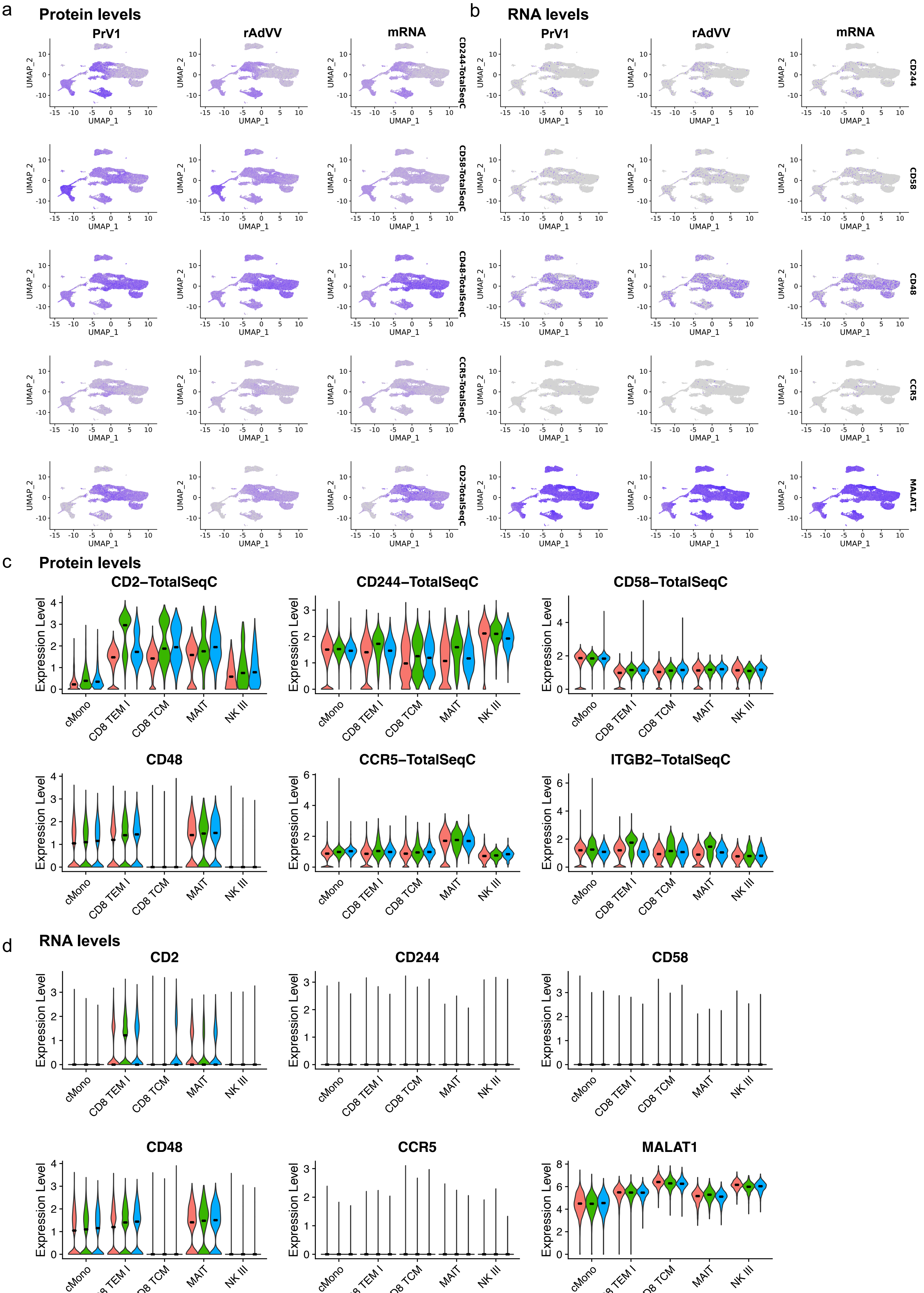
a
Protein levels
PrV1
rAdVV
mRNA
CD244-TotalSeqC
CD58-TotalSeqC
CD48-TotalSeqC
CCR5-TotalSeqC
CD2-TotalSeqC
UMAP_1
UMAP_2
b
RNA levels
PrV1
rAdVV
mRNA
CD244
CD58
CD48
CCR5
MALAT1
c
Protein levels
CD2–TotalSeqC
CD244–TotalSeqC
CD58–TotalSeqC
CD48
CCR5–TotalSeqC
ITGB2–TotalSeqC
Expression Level
cMono
CD8 TEM I
CD8 TCM
MAIT
NK III
d
RNA levels
CD2
CD244
CD58
CD48
CCR5
MALAT1
Expression Level
cMono
CD8 TEM I
CD8 TCM
MAIT
NK III

Suppl. Fig. 11

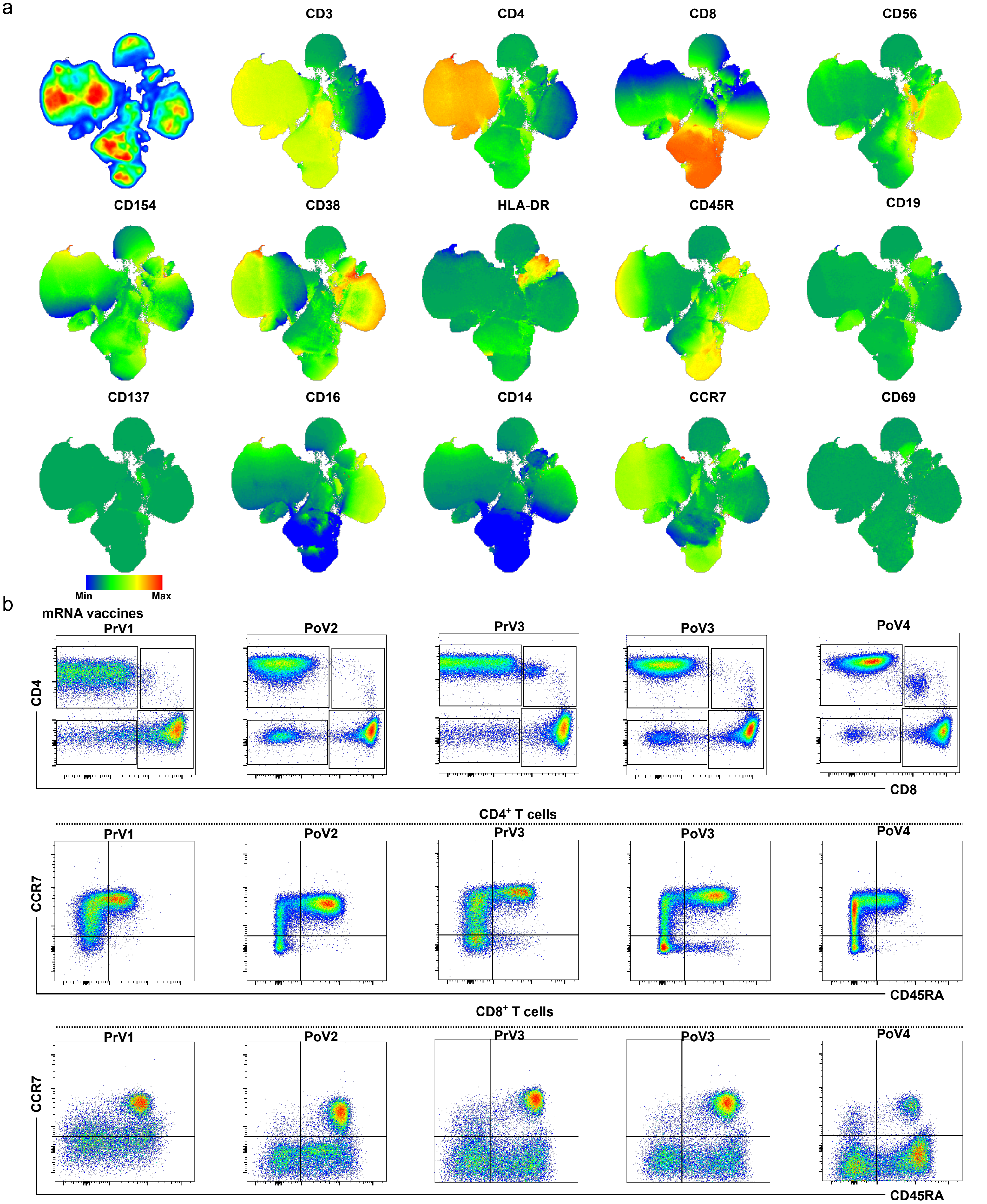
a
CD3
CD4
CD8
CD56
CD154
CD38
HLA-DR
CD45R
CD19
CD137
CD16
CD14
CCR7
CD69
Min
Max
b
mRNA vaccines
PrV1
PoV2
PrV3
PoV3
PoV4
CD4
CD8
CD4+ T cells
CCR7
CD45RA
CD8+ T cells
CCR7
CD45RA

Suppl. Fig. 12

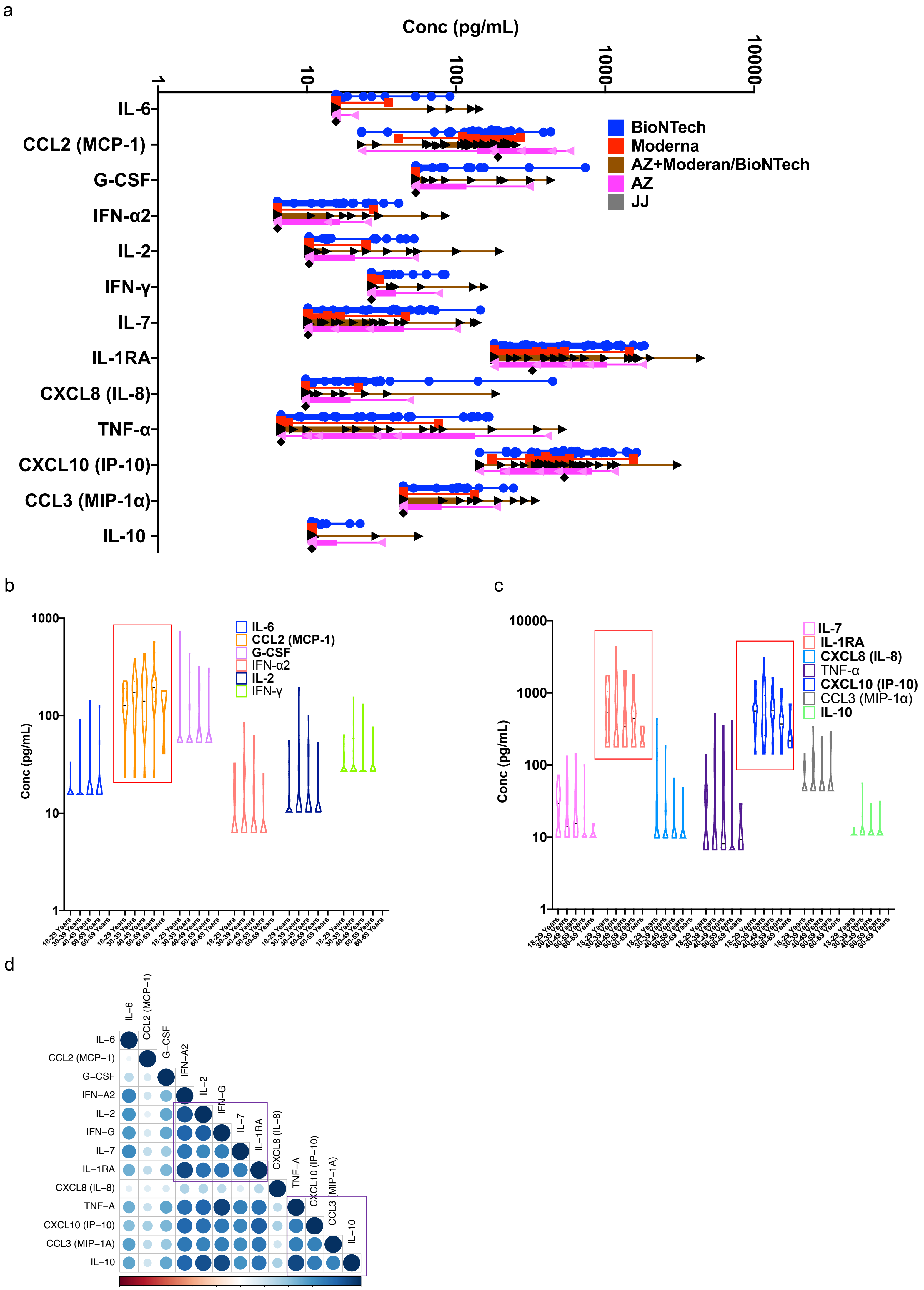
a
Conc (pg/mL)
1
10
100
1000
10000
IL-6
CCL2 (MCP-1)
G-CSF
IFN-α2
IL-2
IFN-γ
IL-7
IL-1RA
CXCL8 (IL-8)
TNF-α
CXCL10 (IP-10)
CCL3 (MIP-1α)
IL-10
BioNTech
Moderna
AZ+Moderan/BioNTech
AZ
JJ
b
Conc (pg/mL)
1000
100
10
1
IL-6
CCL2 (MCP-1)
G-CSF
IFN-α2
IL-2
IFN-γ
18-29 Years
30-39 Years
40-49 Years
50-59 Years
60-69 Years
c
Conc (pg/mL)
10000
1000
100
10
1
IL-7
IL-1RA
CXCL8 (IL-8)
TNF-α
CXCL10 (IP-10)
CCL3 (MIP-1α)
IL-10
18-29 Years
30-39 Years
40-49 Years
50-59 Years
60-69 Years
d
IL–6
CCL2 (MCP–1)
G–CSF
IFN–A2
IL–2
IFN–G
IL–7
IL–1RA
CXCL8 (IL–8)
TNF–A
CXCL10 (IP–10)
CCL3 (MIP–1A)
IL–10
–1 –0.8 –0.6 –0.4 –0.2 0 0.2 0.4 0.6 0.8 1

Suppl. Fig. 13
